# A marker chromosome in psychosis identifies glycine decarboxylase (GLDC) as a novel regulator of neuronal and synaptic function in the hippocampus

**DOI:** 10.1101/2023.05.29.542745

**Authors:** Maltesh Kambali, Yan Li, Petr Unichenko, Jessica Feria Pliego, Rachita Yadav, Jing Liu, Patrick McGuinness, Johanna G. Cobb, Muxiao Wang, Rajasekar Nagarajan, Jinrui Lyu, Vanessa Vongsouthi, Colin J. Jackson, Elif Engin, Joseph T. Coyle, Jeaweon Shin, Michael E. Talkowski, Gregg E. Homanics, Vadim Y. Bolshakov, Christian Henneberger, Uwe Rudolph

## Abstract

The biological significance of a small supernumerary marker chromosome that results in dosage alterations to chromosome 9p24.1, including triplication of the *GLDC* gene encoding glycine decarboxylase, in two patients with psychosis is unclear. In an allelic series of copy number variant mouse models, we identify that triplication of *Gldc* reduces extracellular glycine levels as determined by optical fluorescence resonance energy transfer (FRET) in dentate gyrus (DG) but not in CA1, suppresses long-term potentiation (LTP) in mPP-DG synapses but not in CA3-CA1 synapses, reduces the activity of biochemical pathways implicated in schizophrenia and mitochondrial bioenergetics, and displays deficits in prepulse inhibition, startle habituation, latent inhibition, working memory, sociability and social preference. Our results thus provide a link between a genomic copy number variation, biochemical, cellular and behavioral phenotypes, and further demonstrate that GLDC negatively regulates long-term synaptic plasticity at specific hippocampal synapses, possibly contributing to the development of neuropsychiatric disorders.

## Introduction

Glycine has emerged as an important agonist or modulator of neurotransmitter receptors in the CNS, including NMDA receptors, at which glycine or D-serine are required as co-agonists in addition to the main neurotransmitter glutamate^1, 2^, excitatory eGly (GluN1/GluN3A) receptors^3^ and inhibitory strychnine-sensitive glycine receptors^4^. However, it is largely unknown how glycine levels are regulated in the CNS. Glycine is degraded by the glycine cleavage system (GCS), of which the rate-limiting enzyme is glycine decarboxylase (GLDC)^5^. Inactivating mutations in the human *GLDC* gene have been found in patients with non-ketotic hyperglycemia, a condition with abnormally high levels of glycine in the body resulting in lethargy and hypotonia^6^. In adult brain, GLDC is expressed in astrocytes, but not in neurons^7–9^. While NMDA receptor dysfunction has been postulated to be an important pathophysiological feature of schizophrenia^10,11^, it is unknown whether genetically induced abnormalities in glycine handling may contribute to the pathophysiology of psychosis.

A small supernumerary marker chromosome that segregated with psychosis was discovered in a proband with schizoaffective disorder and his mother with bipolar disorder with psychotic features^12, 13^. This marker chromosome contained genomic material from the small arm of chromosome 9 (9p24.1) with 1.8 Mb length involving 15 genes. Most of the genes were duplicated, however, the *GLDC* gene encoding glycine decarboxylase was triplicated^12, 14^. We decided to use a combination of gene targeting, mutagenesis with CRISPR/Cas-9, and *trans*-allelic recombination *in vivo* to generate an allelic series of the copy number variants (CNVs) in mice to assess the neurobiological significance of this unique CNV and to identify the gene(s) underlying the observed phenotypes using a functional genomic fine-mapping approach. Our present study demonstrates a link between a rare CNV with triplication of GLDC and schizophrenia-like pathophysiological features, and at a fundamental level identifies a physiological role of GLDC as a novel astrocytic negative modulator of long-term synaptic plasticity in the dentate gyrus and cognitive function.

## Results

### Generation and analysis of 9p24.1 CNV (4c9LR) mice

A small supernumerary marker chromosome carrying 15 genes from the 9p24.1 chromosomal region has been identified in two patients with psychosis^12, 14, 15^. To assess the neurobiological significance of this marker chromosome, we generated duplication and deletion alleles for the mouse homologs of the genes on this chromosome (*Rln1*, *Plgrkt*, *Cd274*, *Pdcd1lg2*, *Ric1*, *Emrp1*, *Mlana*, *9930021J03Rik*, *Ranbp6*, *Il33*, *Trpd52l3*, *Uhrf2*, *Gldc*) (for details see Materials and Methods, and **Fig. 1A**). The duplication and deletion alleles were confirmed by comparative genomic hybridization (**Fig. 1B**) and allowed us to breed mice with 1-4 copies of the genomic segment (hereafter called 9LR): deletion (1c9LR), wildtype (2c9RL=WT), duplication (3c9LR) and “triplication” (= homozygous duplication) (4c9LR) mice. Western blots revealed that mice with 1 copy of the 9LR segment expressed significantly less GLDC protein and mice with 4 copies expressed significantly (approximately threefold) more GLDC protein, one of the proteins in the 9LR segment (**Fig. 1C**). We then assessed whether copy number variation leads to changes similar to some of those seen in patients with schizophrenia, such as reduction in dendritic spine density^16^, latent inhibition deficits^17, 18^ and working memory deficits^19^. In Golgi staining the dendritic spine density in dentate gyrus was significantly reduced in 4c9LR mice but not in 1c9LR and 3c9LR mice compared to wildtype (**Fig. 1D**). Mice with 1 and 4 copies of the 9LR segment (1c9LR and 4c9LR) displayed a deficit in latent inhibition to conditioned freezing, whereas in mice with 2 (WT) and 3 copies of 9LR segment latent inhibition was present (**Fig. 1E**), indicating that latent inhibition is sensitive to copy number of the 9LR segment. In a water T maze paradigm in which rewarded alternation was tested, introduction of a 45 second delay between forced and choice trials reduced the percentage of correct responses in mice with 3 and 4 copies of 9LR (3c9LR and 4c9LR mice) (**Fig. 1F**), indicating that an increase in the copy number of the 9LR segment results in a working memory deficit. 4c9LR mice showed no difference in hippocampal-independent delay conditioning (**Extended Data Fig. 1A**), but in hippocampal-dependent trace fear conditioning (TFC) (**Extended Data Fig. 1B**) and contextual conditioning (**Extended Data Fig. 1C**), they displayed increased freezing. As the patients with the 9p24.1 marker chromosome had symptoms of both psychotic and mood disorders, we also assessed depression-related behavioral changes in these mice. A reduction in sucrose preference in 4c9LR mice indicated anhedonia (**Extended Data Fig. 1D).** Novelty-suppressed feeding (**Extended Data Fig. 1E**), motor activity across the light/dark cycle (**Extended Data Fig. 1F**), and anxiety-like behavior in the light/dark test (**Extended Data Fig. 1G**) and in the elevated plus maze (**Extended Data Fig. 1H**) were unaltered. Motor activity in a familiar open field test was increased in 1c9LR and 4c9LR mice after group-housing but not after single housing (**Extended Data Fig. 1I**). In the forced swim test, immobility was increased in 3c9LR and 4c9LR mice after 6 weeks of single housing isolation stress but not after group housing (**Extended Data Fig. 1J**), indicating that social isolation stress may play a role.

**Figure 1.**
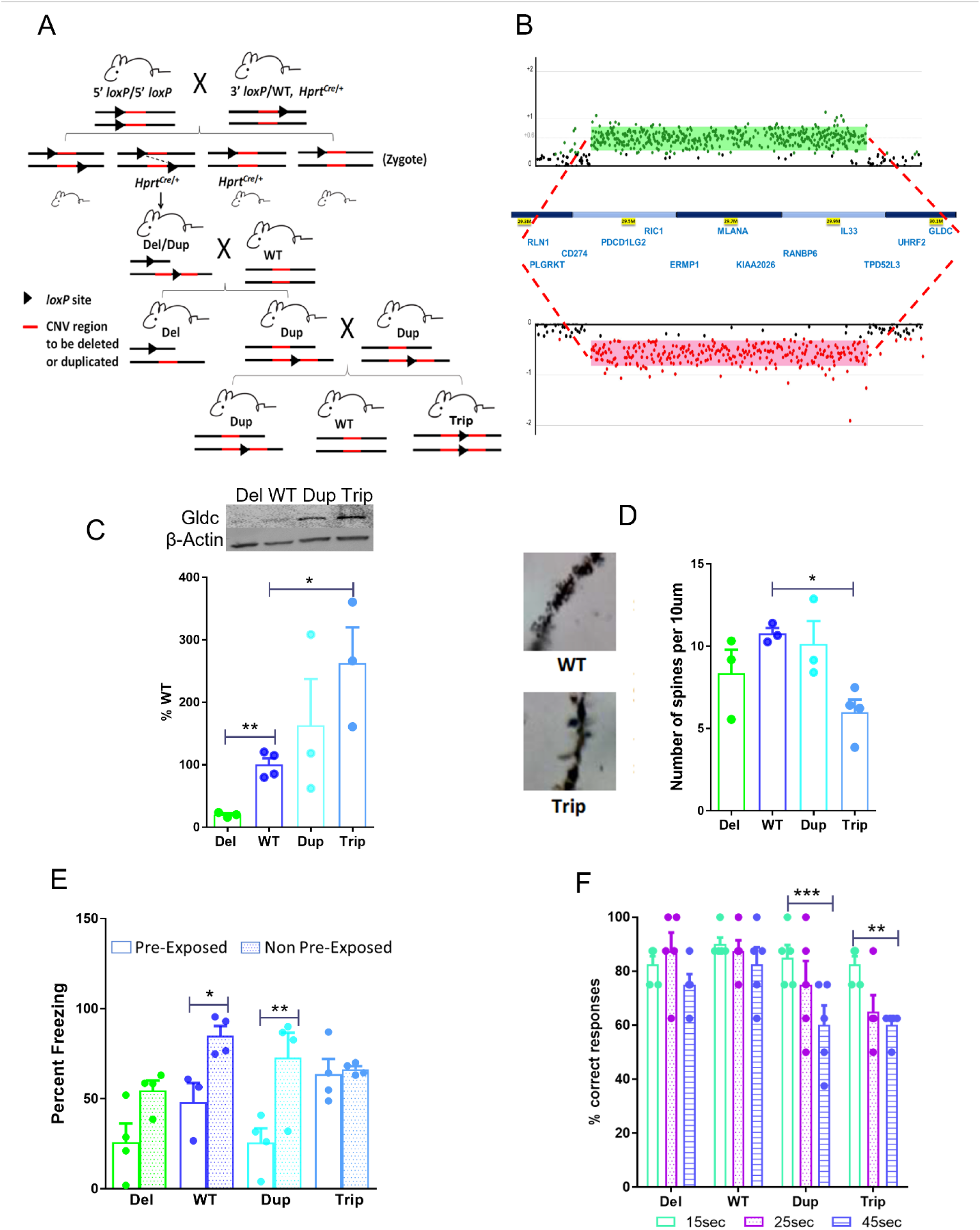
Schematic representation of the generation of the 9p24.1 CNV mice. A. Generation of the 9p24.1 deletion (Del), duplication (Dup) and triplication (Trip) alleles by selective breeding with targeted *lox*P sites in *trans* and a Hprt-Cre transgene, via *trans*-allelic recombination. B. Comparative genomic hybridization for the duplication (green) and deletion (red) alleles using an Agilent 1M array. C. Representative western blots and quantification of GLDC protein expression in hippocampus of 9p24.1 deletion (Del=1c9LR), duplication (Dup=3c9LR) and triplication (Trip=4c9LR) mice compared to wildtype (WT=2c9LR) (expressed as percentage of wildtype, unpaired two-sided Student’s t-test; 4c9LR vs WT; t(5)=3.269, *p<0.05, 1c9LR vs WT; t(5)=6.540, **p<0.01, n=3-4 for each group). D. Golgi staining was performed to determine dendritic spine density in dentate gyrus, suggesting a reduced density in triplication (Trip) mice (n=3-7 dendrites counted in 3 mice per genotype, One-way ANOVA followed by Bonferroni post hoc test, F(3,12)=4.624; p=0.032; 4c9LR vs WT; t=3.341, *p<0.05). E. Latent inhibition to conditioned freezing revealed absence of latent inhibition in 9p24.1 deletion (Del) and triplication (Trip) mice, i.e.,, the percentage time spent freezing during presentation of the tone was not different in mice that were pre-exposed to the tone [30 tones (20 sec, 70 dB, 2,800 Hz) with 30 sec intervals] versus non-pre-exposed mice, whereas pre-exposed WT and 9p24.1 duplication (Dup) mice were freezing less than non-pre-exposed mice (n=4-5 per group, One-way ANOVA followed by Bonferroni’s post-hoc test, F(7,23)=6.344; p=0.0003; WT-pre vs non-pre exposed; t=2.867, *p<0.05, Dup-pre vs non-pre exposed; t= 3.953, **p<0.01). F. Water T-maze non-matching to place task to test working memory. Mice were trained to alternate the arms visited in a water T-maze with typically >70% accuracy. On test days, different delay intervals between forced trials and choice trials were applied (15 s, 25 s and 45 s) (n=5-6/group, Two-Way ANOVA revealed main effect of genotype (F(3,16)= 3.882, p<0.05) and of delay (F(2,32)= 13.48, p<0.001) followed by Bonferroni posthoc test revealed at 45 sec vs 15sec; Dup, t= 4.126, ***p<0.001, and Trip, t= 3.713, **p<0.01).

### RNA sequencing of mice with 1-4 copies of the marker chromosome genes

RNA sequencing revealed that expression of the genes in the 9p24.1 cytoband correlated with CNV genotype in both hippocampus (top panel, HPC) and prefrontal cortex (bottom panel, mPFC) **(Extended Data Fig. 2A)**. PCA scatter plots show tissue-level variation as strongest among known batch variables **(Extended Data Fig. 2B, C, D)**. Differentially expressed genes (DEGs) were also correlated with genotype as the largest number of DEGs were observed in triplication mice in the HPC and in duplication mice in the mPFC **(Fig. 2A-D, Supplementary tables 1a-1f** and **Extended Data Fig. 2E-H)**. KEGG-pathway analysis revealed long-term potentiation and dopaminergic synapse as downregulated pathways, while ECM-receptor interaction and cell adhesion pathways were upregulated pathways in HPC of triplication mice (**Fig. 2E, F**). For other genotypes, see **Supplementary tables 2a-2r**. Significant enrichment was observed for DEGs from 9p24.1 CNVs with rare coding variants among genes linked to autism and neurodevelopmental disorders (NDD) from a recent large-scale sequencing study (**Fig. 2G and supplementary table 3**)^20^. Furthermore, *Pyroxd2* (pyridine nucleotide-disulphide oxidoreductase domain 2), a gene encoding a protein involved in mitochondrial-respiration^21^, was upregulated in all mutant genotypes in both the HPC and the mPFC **(Fig. 2H, I)**. Mutations or deletion of PYROXD2 in humans leads to increased oxidative-stress and mitochondrial dysfunction^22^. Furthermore, the expression of *Arl3* (ADP-ribosylation factor-like GTPase 3), which has been associated with schizophrenia by common variant effects from a genome wide association study (GWAS)^23^, was reduced in duplication and triplication genotypes in both HPC and mPFC **(Fig. 2J, K)**.

**Figure 2.**
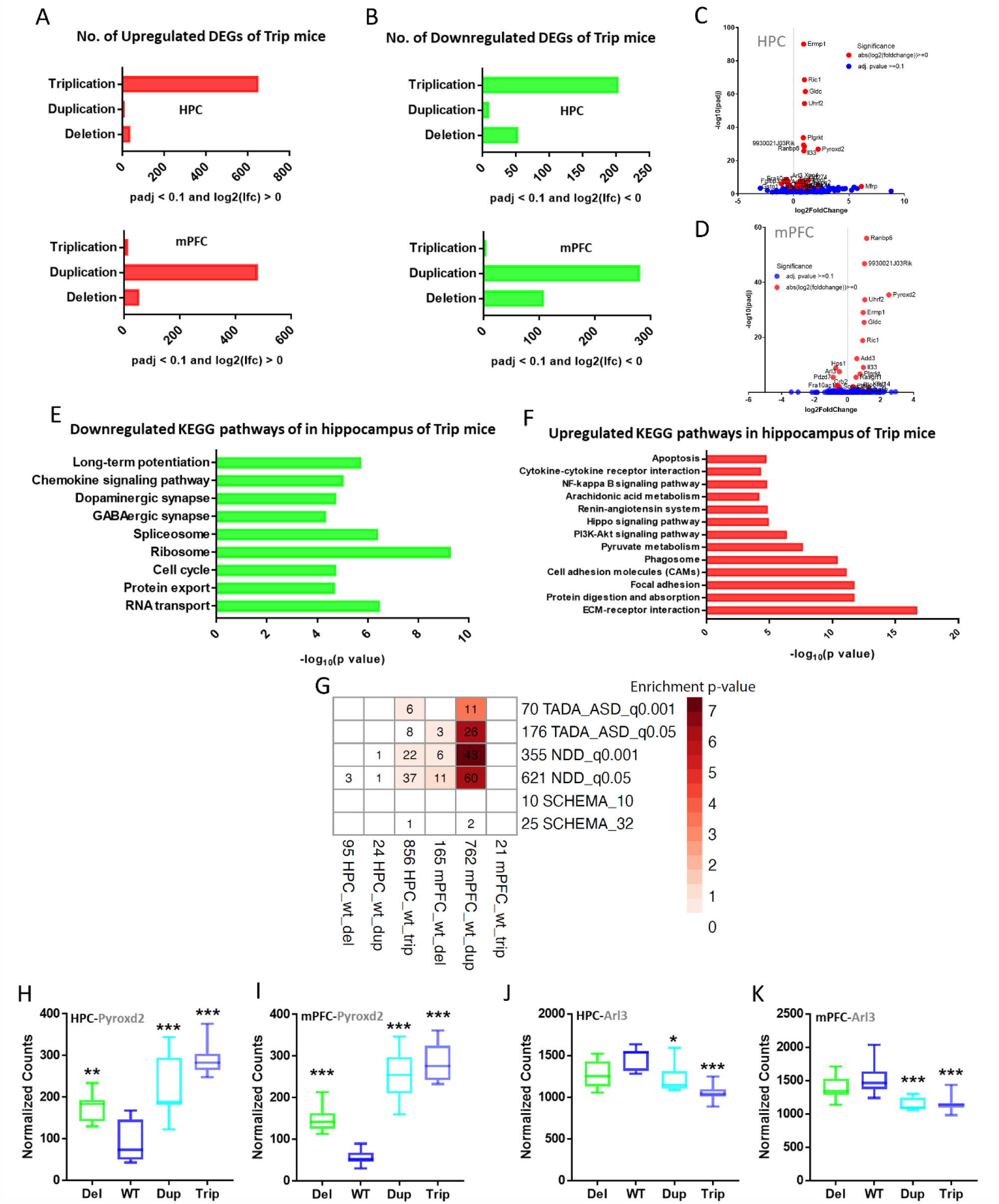
Gene expression changes in hippocampal and prefrontal cortex with KEGG enrichment pathways, and associated enrichment with TADA-ASD and NDD gene sets. A, B. The bar graphs show the number of differentially expressed genes (DEG) upregulated (red) and down regulated (green) in hippocampus and prefrontal cortex of triplication mice with the Adj. P value <0.1. The volcano plots show the distribution of log2(fold change) and associated negative log_10_(padj) for hippocampus (C) and prefrontal cortex (D) of triplication (Trip) mice as compared to wild type, with top 30 DEGs highlighted in red. The KEGG pathway classification of genes into enriched pathway terms (E. green, down-regulated pathways; F. red, up-regulated pathways) in hippocampus of triplication mice. The KEGG pathway enrichment cut-offs of significance from FDR < 0.1 to p-value <0.05 was criteria. G. DEG enrichment for gene sets previously published with associations with neurological phenotype; Autism Spectrum Disorder (ASD): Transmission and de novo association with autism spectrum disorder; Neurodevelopmental Disorder-associated genes (NDD): Neurodevelopmental Disorder-associated genes^20^. The top two among the DEGs are, Pyroxd2 showed upregulation in all the mutant genotypes in both H. Hippocampus, and I. prefrontal cortex, and Arl3 showed down- regulation in duplication and triplication genotypes at both J. Hippocampus and K. prefrontal cortex. The data presented as 10-90 percentile, (Hippocampus: n=7-8/group, prefrontal cortex: n=8/group), One-Way ANOVA followed by Bonferroni test, *p<0.05, **p<0.01, ***p<0.001.

### Genomic fine mapping of gene(s) underlying the observed behavioral phenotype

To identify the gene(s) underlying the observed phenotype, we generated mice with two smaller, complementary CNVs, one including only *Gldc* (3c9R and 4c9R), encoding a glycine-degrading enzyme which might modulate availability of glycine, a co-agonist at the NMDA receptor, and the other one only including the remaining genes of the 9LR segment (3c9L and 4c9L, for details see Materials and Methods and **Fig. 3A**). In 4c9R mice, expression of GLDC was increased approximately two-fold, similar to 4c9LR mice, while in 4c9L mice GLDC expression was unchanged **(Fig. 3B)**, in line with the *Gldc* copy number.

**Figure 3.**
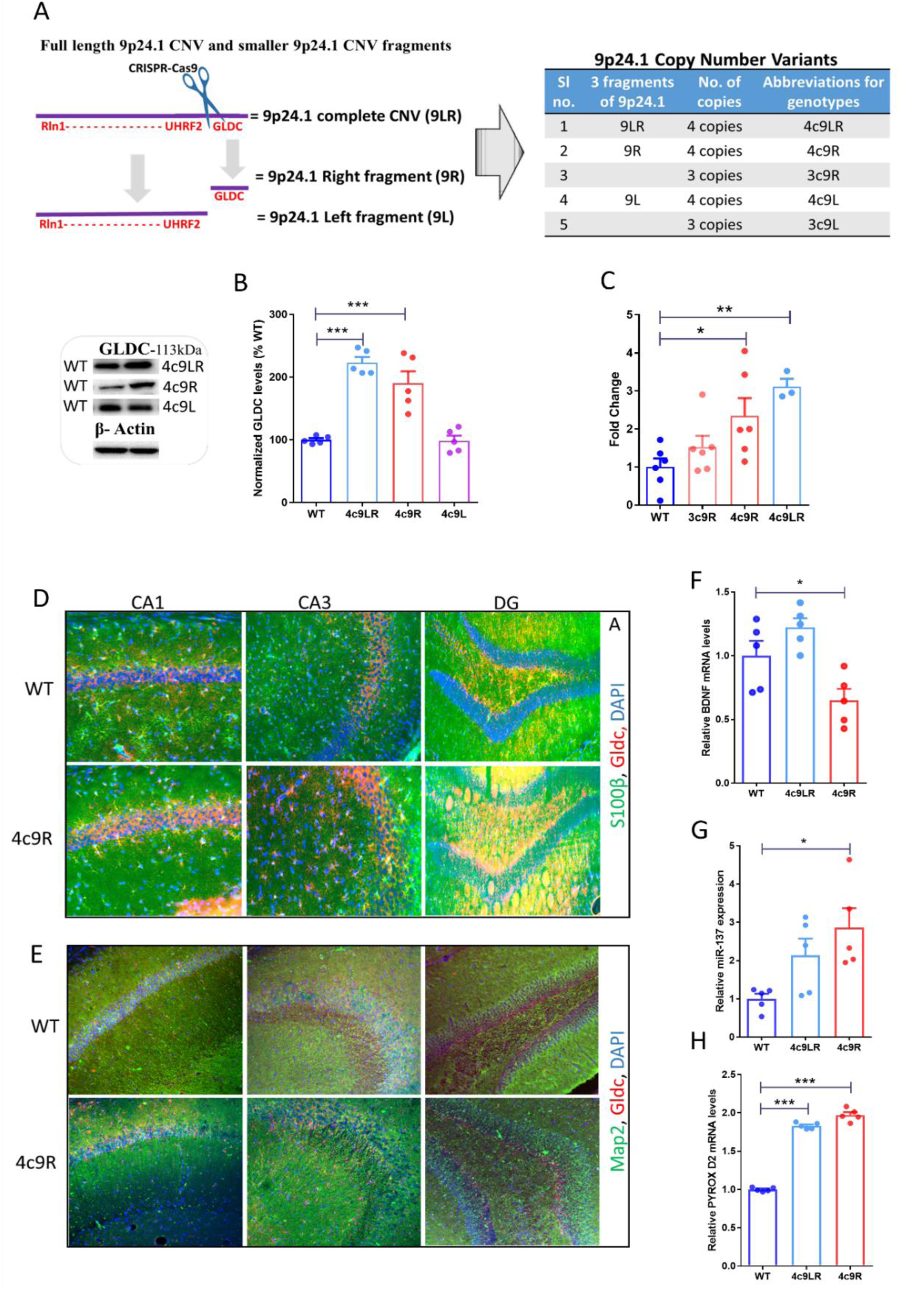
A. Generation and characterization of shorter complementary CNVs from the 9p24.1 CNV (9LR) using CRISPR/Cas-9 genome editing. A loxP site was placed between the *Uhrf2* and *Gldc* genes using CRISPR/Cas-9, and *trans*-allelic recombination was used as described in Methods to generate duplication alleles of *Gldc* alone (9R fragment; duplication=3c9R, triplication=4c9R) or of all the genes in the 9LR genomic segments (9L fragment; duplication=3cL, triplication=4cL). B. Representative western blots and quantification of GLDC protein in hippocampus of mice-with 4 copies of GLDC alone (4c9R), 4 copies of all other 14 genes of the 9p24.1 CNV (4c9L) and 4 copies of the entire 9p24.1 CNV(4c9LR) compared to wildtype (WT) (n=5 for each group, One-way ANOVA followed by Bonferroni test, F(3,37)=18.77; ***p<0.001, WT vs 4c9LR (t=6.07; ***p<0.001), WT vs 4c9R (t=5.24;***p<0.001). C. GLDC enzyme activity in hippocampal tissues of wild type (WT), 4c9LR, 4c9R and 3c9R (n=3-6, One-way ANOVA followed by Bonferroni test, F(3,16)=13.53; ***p=0.0001), WT vs 4c9LR (t=5.827; **p<0.01), WT vs 4c9R (t=3.366; *p<0.05). D, E. Qualitative fluorescence microscopy images of GLDC protein colocalization with S100beta (D) or Map2 (E) in CA1, CA3 and dentate gyrus regions of hippocampus, respectively, in wildtype (WT) and 4c9R mice. F. BDNF mRNA expression is reduced in 4c9R mice compared to WT (One-way ANOVA followed by Bonferroni test; F(2,17)=7.669; **p<0.01), WT vs 4c9R (t=2.729; *p<0.05). G. miR137 mRNA expression is increased in 4c9R mice compared to WT (One-way ANOVA followed by Bonferroni test, F(2,12)=5.683; *p<0.05, WT vs 4c9R (t=3.343, *p<0.05). H. Pyroxd2 mRNA expression is increased in hippocampus of 4c9LR and 4c9R mice (One-way ANOVA followed by Bonferroni test, F(2,12)=468.4; ***p<0.0001, 4c9LR vs WT (t=24.20; ***p<0.001) and WT vs 4c9R (t=28.33; ***p<0.001). Data are mean ± SEM, (n=5).

### Increased GLDC enzymatic activity and increased expression in astrocytes of hippocampus of 4c9R mice

GLDC is a rate limiting enzyme of the glycine cleavage system (GCS)^24^. Both 4c9R and 4c9LR mice, containing 4 copies of the *Gldc* gene, display higher glycine cleavage system (GCS) activity than wild-type mice as seen in hippocampus (**Fig. 3C**), which we focused on as it has been linked to psychosis^25^ and since glycine is known to modulate excitatory neurotransmission in the dentate gyrus^1^. Previous reports indicate that GLDC is expressed in astrocytes but not in neurons^8, 9^. In order to assess whether this also holds true in mice with additional copies of the *Gldc* gene and increased expression of the GLDC protein, we performed immunofluorescence staining to assess the localization of GLDC. Qualitative images in CA1, CA3, and DG show a prominent colocalization of GLDC with astrocytic marker S100b in both wildtype and 4c9R genotypes (**Fig. 3D**) compared to the colocalization with the neuronal marker Map2 (**Fig. 3E**), and staining for GLDC may be increased in 4c9R mice compared to WT. These findings suggest that the increased expression of GLDC in 4c9R mice does not alter the cellular distribution.

### BNDF*, Pyroxd2* mRNA expression and regulatory microRNA dysregulation in 4c9R mice

We next examined expression of genes that have been linked to schizophrenia. BDNF has been found to be reduced in the brain of patients with schizophrenia^26^. Using qRT-PCR, we found that hippocampal BDNF mRNA was reduced in 4c9R mice **(Fig. 3F)**. MicroRNA miR-137 has been reported to be associated with an increased risk of developing schizophrenia in GWAS studies^23^ and its dysregulation has been shown to impair synaptic plasticity in hippocampus^27^. miR-137 expression was increased in hippocampus of 4c9R mice but not in 4c9LR mice **(Fig. 3G)**, providing another link between our 4c9R mice and schizophrenia risk genes identified by GWAS. In line with previous transcriptomics results (**Fig. 2H**), qRT-PCR showed an approximately two-fold increase in *Pyroxd2* mRNA expression in 4c9RL and 4c9R mice (**Fig. 3H**). Thus, while the triplication of the *Gldc* gene alone was necessary and sufficient to cause changes in the expression of some schizophrenia-related genes, others could be altered by the triplication of either part of the whole CNV region.

### Increased GLDC expression impaired synaptic plasticity-related signaling pathways

We next examined BDNF protein levels and the activity of downstream pathways in 4c9R mice compared to WT. In hippocampal synaptoneurosomal fractions we detected a significantly reduced amount of BDNF protein (**Fig. 4A)**. The ratios of phosphorylated, i.e., activated forms of Akt (pAkt-S473 and pAkt-T308) to total Akt were reduced (**Fig. 4B** and **C**), with no change in total Akt (**Fig. 4D**). In addition, the ratio of phosphorylated mammalian target of rapamycin (p-mTOR-S2448) to total mTOR, which is phosphorylated by Akt, was reduced (**Fig. 4E**), with no changes in total mTOR (**Fig. 4H**). When examining the downstream target of mTOR signaling, P70S6K, we did not find any changes in the ratio of phosphorylated P-P70S6K to total P70S6K or in total P70S6K, a protein involved in ribosome biogenesis and protein synthesis^28^ (**Fig. 4F** and **I**). However, when we analyzed a downstream target of Akt, CREB, which regulates gene expression in activity-dependent synaptic plasticity, the ratio of phosphorylated CREB (P-CREB) to total CREB protein was reduced in the hippocampus of 4c9R mice, with no change in total CREB. (**Fig. 4G** and **J**). Thus, there was a dampening of both transcriptional regulation through the transcription factor CREB and translational regulation through the protein kinase mTOR activity downstream of the BDNF-Akt pathway in 4c9R mice, similar to disruptions in BDNF signaling in schizophrenia. These findings demonstrate multilevel effects of increased GLDC expression at the biochemical level.

**Figure 4.**
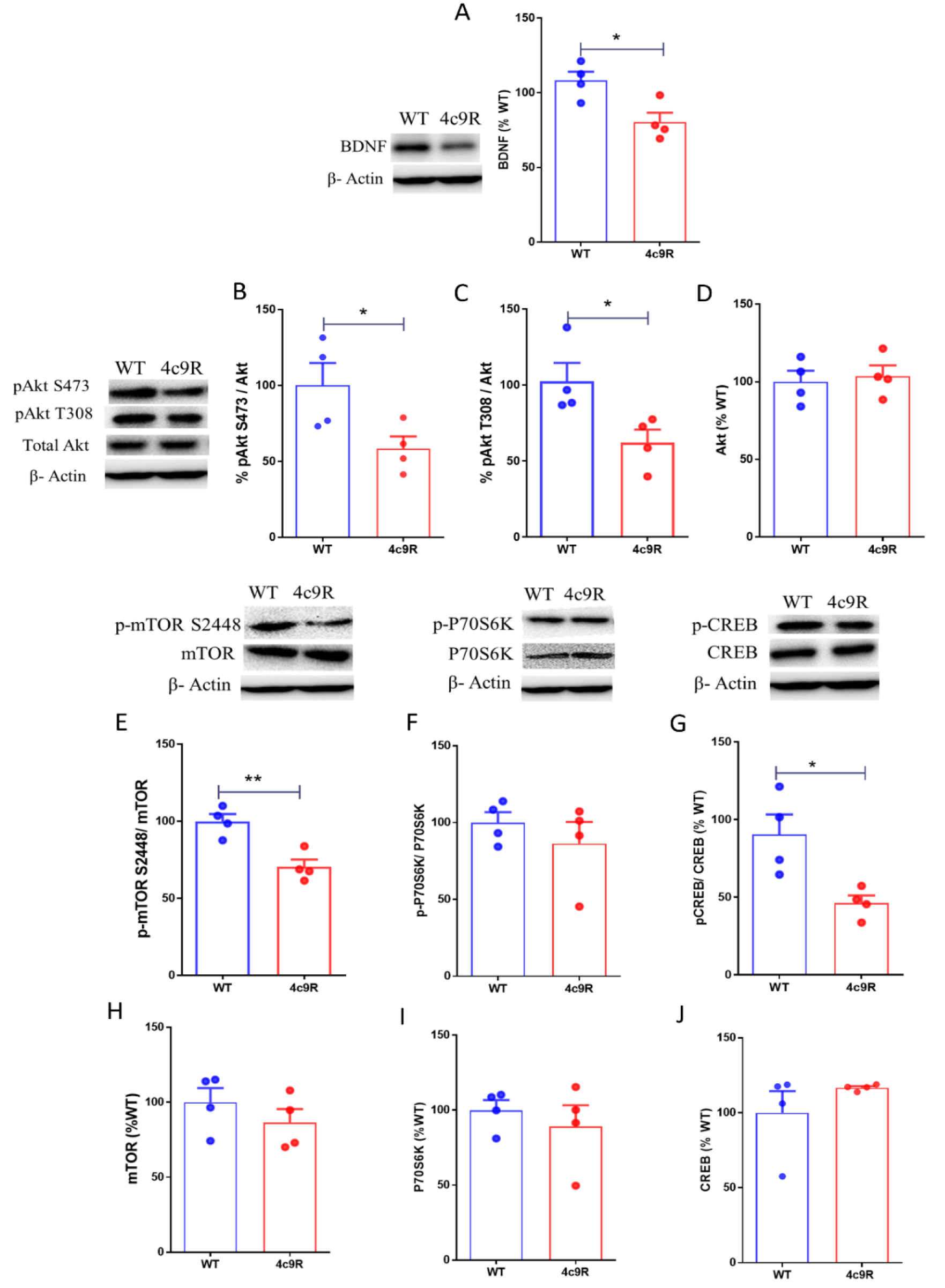
Akt/mTOR signaling and expression of BDNF in synaptoneurosomal fraction of the hippocampus in 4c9R mice. Western blot images and quantification of proteins: A. BDNF (4c9R vs Wildtype; t(6)= 2.208; *p=0.044), B. S473 p-Akt/ Akt (4c9R vs Wildtype; t(6)= 2.743; *p=0.0336), C. T308 p-Akt/ Akt (4c9R vs Wildtype; t(6)= 2.488; *p=0.047), D. Total Akt, E. S2448 p-mTOR/ mTOR (4c9R vs Wildtype; t(6)= 4.413; **p=0.0045), F. p-P70S6K/ P70S6K and G. p- CREB/ CREB (4c9R vs Wildtype; t(6)= 3.195; *p=0.0187), H. Total mTOR, I. Total P70S6K, J. Total CREB. All data presented as mean ± SEM, (n=4, unpaired two-sided Student t-test).

### Reduced extracellular glycine levels in dentate gyrus but not in CA1 in 4c9R mice

Next, we tested the hypothesis that the increased GLDC activity in 4c9R mice **(Fig. 3C**) leads to reduced extracellular glycine levels in the hippocampus, resulting in NMDA receptor dysfunction. Extracellular glycine levels were assessed by anchoring the fluorescent glycine sensor GlyFS, via a biotin-streptavidin linker, in the extracellular space of the CA1 (st r) or DG regions (st m) of biotinylated acute hippocampal slices^29^. The GlyFS fluorescence intensity ratio (*R*0/*R*MAX) was used to estimate the resting glycine levels. In the DG a significant decrease in glycine levels was observed in 4c9R mice compared to WT, whereas in the CA1 subregion there was no difference (**Fig. 5A**). Furthermore, we characterized the activity-dependent mechanisms that control extracellular glycine levels. Because glycine is involved in NMDAR-dependent synaptic plasticity^1, 31^, we focused on plasticity-inducing stimuli. In both the 4c9R mice and WT mice, high-frequency stimulation of the Schaffer collateral CA3-CA1 synapses (**Extended Data Fig. 3A, B**) but not of the performant pathway-DG synapses (**Extended Data Fig. 3C, D**) resulted in increased GlyFS-reported extracellular glycine levels.

**Figure 5.**
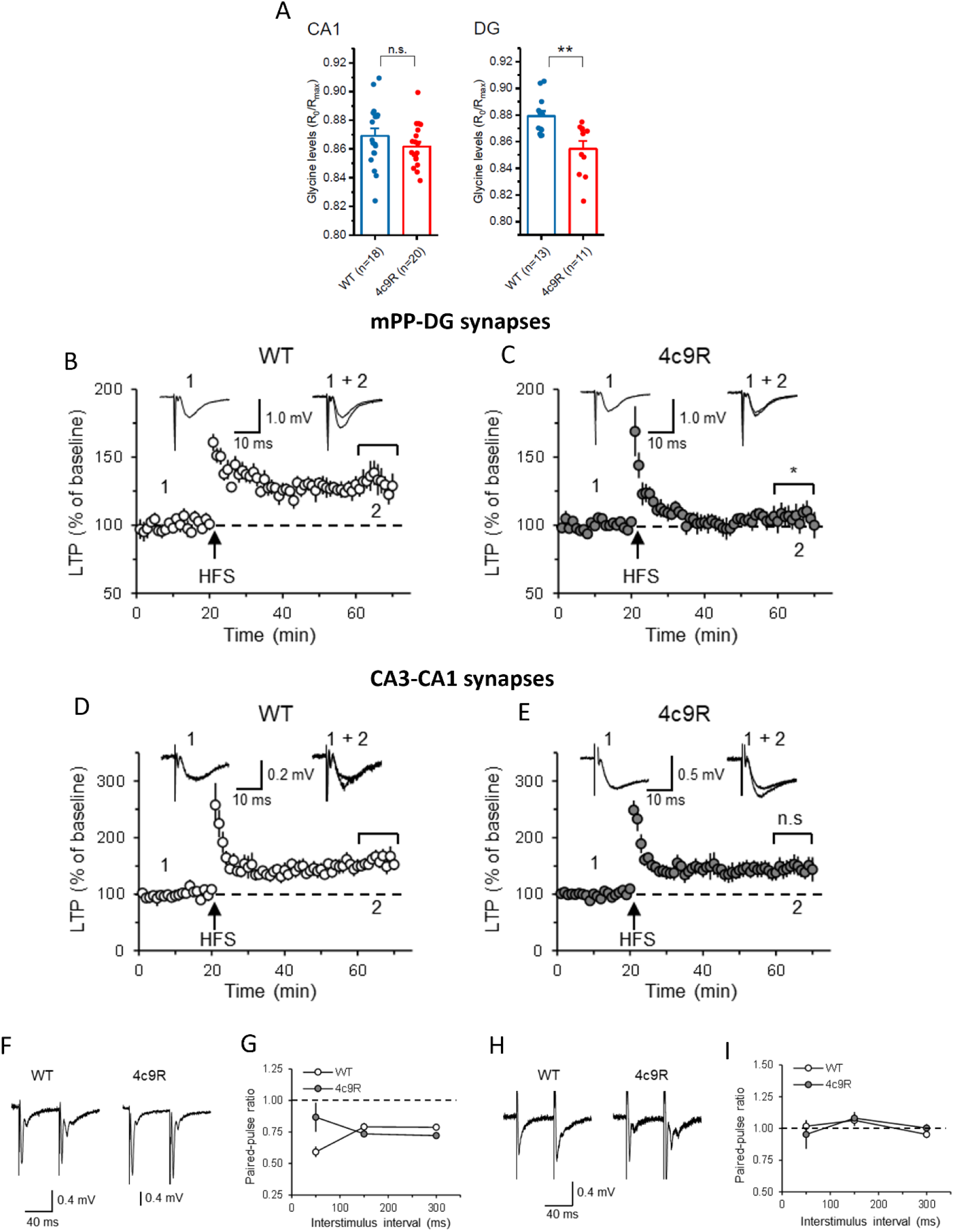
Extracellular synaptic glycine levels measured using optical glycine FRET sensor (GlyFS) in the CA1 and dentate gyrus, and Long-term potentiation in CA3-CA1 synapses and mPP-dentate gyrus synapses of 4 copy GLDC mice. A. Basal extracellular glycine levels are decreased in dentate gyrus but not in CA1 as measured by GlyFS R0/Rmax ratio from 4c9R mice compared to wildtype (unpaired two-sided Student’s t-test, t(22) = 3.59, **p<0.01 at DG, WT n = 13, 4c9R n = 11). LTP is suppressed at the mPP-DG synapses (B and C) but not at the CA3-CA1 synapses (D and E) in 4c9R mice. Summary of LTP experiments from four wildtype mice (n = 5 slices) and four 4c9R mice (n = 8 slices). LTP was induced by a 1-s train of 100 Hz stimulation. Insets in B and C are the averages of 40 fEPSPs recorded before (1) and 20 fEPSPs recorded 40 min after (2) the induction (at arrow) of LTP in slices from WT and 4c9R mice. Asterisk (*) indicates significant differences between the two groups (*P < 0.05, unpaired two-sided student t-test). All values represent the mean ± SEM. But LTP is not suppressed in CA3-CA1 synapses in 4c9R mice (D and E). Summary of LTP experiments from four wildtype mice (n = 8 slices) and four 4c9R mice (n = 7 slices). Insets in D and E are the averages of 40 fEPSPs recorded before (1) and 20 fEPSPs recorded 40 min after (2) the induction (at arrow) of LTP in slices from WT and 4c9R mice. Examples of fEPSPs at the (F) mPP-DG synapses and (H) CA3-CA1 synapses evoked with paired stimuli (interstimulus intervals 50-ms) in slices from WT and 4c9R mice. Averaged paired-pulse ratio values at the (G) mPP-DG synapses and (I) CA3-CA1 synapses in slices from four WT (n = 7 slices) and four 4c9R (n = 7 slices) mice at interstimulus intervals 50, 150, and 300 ms. Paired-pulse ratio was calculated as the ratio of the rising slope of the second fEPSP to the first fEPSP.

Overall, the experiments using the recently developed GlyFS revealed that an increased copy number of *Gldc* which we have shown to increase the activity of the GLDC enzyme results in a reduction of extracellular glycine levels in the DG but not in CA1, and that – independently of the *Gldc* copy number – extracellular glycine levels in CA1 and DG display differential sensitivities to HFS. As it has been shown previously that glycine, and not D-serine is the major NMDA receptor co-agonist in the DG (while the opposite is true for CA1)^1^, the observed decreased extracellular glycine levels in the DG could translate into selective NMDA receptor hypofunction in the DG.

### Diminished Long-Term Potentiation at the Perforant Pathway to DG Synapses but not at CA3-CA1 synapses in 4c9R mice

To assess whether NMDAR hypofunction in 4c9R mice results in a reduction of basal neurotransmission and/or of NMDAR-dependent long-term synaptic potentiation (LTP), we stimulated the medial-perforant-pathway (mPP) or Schaffer-collaterals-fibers pathway (SCF) and recorded field excitatory postsynaptic potentials (fEPSPs) in the DG (st-m) or CA1 (st-r) regions of hippocampal slices from WT and 4c9R mice respectively. The initial potentiation after the LTP-inducing high-frequency stimulation was similar in both pathways between WT and 4c9R mice and likely reflects post-tetanic potentiation or short-term potentiation, forms of synaptic plasticity independent of NMDAR activation^32^. Strikingly, the magnitude of LTP was significantly reduced in mPP-DG pathway from 4c9R mice compared to WT (**Fig. 5B, C**) (unpaired two-sided student t-test, p<0.05). However, the magnitude of LTP was not altered in CA3-CA1 synapses (**Fig. 5D, E**). The paired pulse ratio at mPP to DG (**Fig. 5F, G)** and CA3-CA1 synapses (**Fig. 5H, I**) remained unchanged in slices from 4c9R mice. Overall, our electrophysiological studies revealed that NMDA receptor-dependent synaptic plasticity was significantly reduced in 4c9R mice, which could be explained by our finding that extracellular glycine concentrations in the DG are reduced (**Fig. 5A**).

### Reduced dendritic spine density and differential expression of GLDC and H3K4me3 histone methylation in dentate gyrus and CA1

A reduced spine density is one of the most consistently observed neuropathological alterations in schizophrenia^33, 34^. The percentage of mature mushroom spines, which have been linked to memory functions^35, 36^, was significantly reduced in 4c9R compared to wildtype (**Fig. 6A**), and the percentage of stubby type spines increased (**Fig. 6B**). The percentage of thin spines was not altered (**Fig. 6C**). The total spine density (protrusions) was reduced in 4c9R mice (**Fig. 6D**) compared to wildtype. In line with this observation, the microRNA mir132, a regulator of dendritic spine structure^37^ which is downregulated in peripheral blood of patients with schizophrenia^38^, was reduced in hippocampus of both genotypes with 4 copies of GLDC, 4c9R and 4c9LR **(Fig. 6E)**.

**Figure 6.**
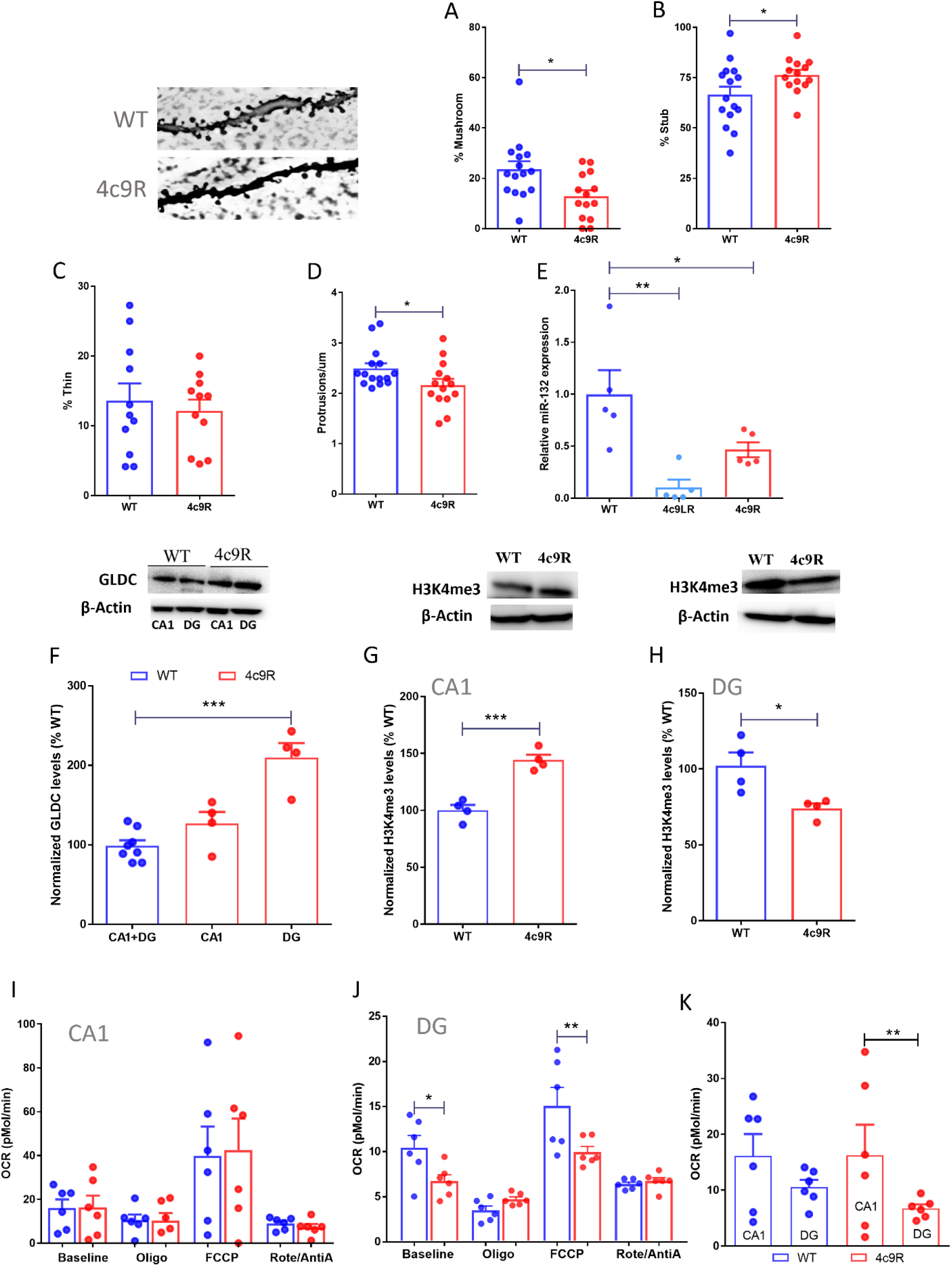
Effect of 4 copies of *Gldc* on dendritic spine morphology, mitochondrial bioenergetics and histone methylation in dentate gyrus and CA1. Images of dendritic branches with spines from dentate gyrus from WT and 4c9R mice. Dendritic spine quantification shows a reduced percentage mushroom type spines (A) (n= 14-15 spines, unpaired two sided Student t-test, t(27)=2.639, *p<0.05), and an increased percentage of stubby type spines (B) (n= 14-15 spines, unpaired two sided Student t-test, t(27)=2.058, *p<0.05). C. There was no change in the percentage of thin spines. D. The total number of spines (protrusions/um) on secondary dendritic branches is reduced in dentate gyrus of 4c9R mice compared to WT (four mice each group, n= 14-15 spines, unpaired two sided Student t-test, t(27)=2.094, *p<0.05). E. Lower amounts of microRNA miR132 in hippocampus of 4c9Rmice and 4c9LR mice (n=5, One-way ANOVA followed by Bonferroni test, F(2,12)=9.524; **p<0.01; WT vs 4c9LR (t=4.337; **p<0.01) and WT vs 4c9R (t=2.588; *p<0.05)). F-H. Hippocampal sub-regional expression of GLDC and H3K4me3 proteins in CA1 and dentate gyrus of 9c4R mice. F. GLDC expression in CA1 and dentate gyrus in 4c9R mice compared to wildtype (WT) (n=4, 3 male+ 1 female mice per group, One-way ANOVA followed by Bonferroni test, F(2,13)=22.73; ***p<0.0001, DG; t=6.718, ***p<0.001). G, H. The histone methylation marker H3K4me3 is increased in CA1 of 4c9R mice (G) (4c9R vs WT, unpaired two-sided student t-test; t(6)=6.687, ***p<0.001) and decreased in dentate gyrus of 4c9R mice (H) (WT vs 4c9R, unpaired two-sided student t-test; t(6)=3.055; *p<0.05, n=4, 3 male+ 1 female mice per group). I-K. Seahorse mitochondrial stress test. The OCR measurements show the characteristic responses to mitochondrial inhibitors and the uncoupler FCCP. Effect of 4 copies of *Gldc* on mitochondrial oxygen consumption rate (OCR) in tissue punches from hippocampal subregions. I. No change in CA1, J. Decreased basal OCR in DG (One-way ANOVA followed by Bonferroni test, F(7,16)=18.54, ***p<0.0001; 4c9R vs WT, t=2.962, *p<0.05), and reduced FCCP response (One-way ANOVA followed by Bonferroni test; F(7,16)=18.54, ***p<0.0001; 4c9R vs WT, t=4.209, **p<0.01). K. CA1 vs DG basal OCR (Two-way ANOVA revealed main effect of 4 copies of GLDC (F (3, 48) = 6.965, ***p<0.001) followed by Bonferroni test, DG vs CA1 basal OCR in 4c9R, t=5.426, **p<0.01). For all groups, 4 males and 2 female mice (n=6) were used. All data are presented as mean ± SEM.

In view of findings that the additional copies of *Gldc* affect extracellular glycine levels differentially in DG and CA1 (**Fig. 5A**), we assessed whether there are also other hippocampal sub-regional differences. The 4c9R mice displayed significantly higher GLDC expression in DG but not in CA1 (**Fig. 6F**), which could explain why extracellular glycine was reduced in 4c9R mice in DG but not CA1. BDNF protein expression was decreased in 4c9R mice in both CA1, and DG compared to WT (**Extended Data Fig. 4E, F**). In view of a report that in pluripotent stem cells the activated glycine cleavage system catabolizes glycine to fuel H3K4me3 trimethylation^24^, we investigated whether additional copies of *Gldc* would increase H3K4me3 trimethylation in the hippocampus. The H3K4me3 trimethylation was significantly increased in CA1 but significantly decreased in DG in 4c9R mice (**Fig. 6G, H**). Decreased levels of H3K4me3 associated with decreased GAD1 expression have been reported in prefrontal cortex from schizophrenia patients^39^, indicating that altered levels of H3K4me3 may play a role in the pathophysiology of schizophrenia. The functional significance of the differential modulation of H3K4me3 in DG and CA1 is currently unclear.

### Attenuation of mitochondrial bioenergetics in dentate gyrus of 4c9R mice

Cerebral organoids derived from patients with schizophrenia display reduced basal oxygen consumption rates (OCR)^40^. We found that in CA1 basal respiration, respiration after inhibition of ATP-linked respiration by oligomycin and FCCP (carbonyl cyanide-p-trifluoromethoxyphenyl-hydrazon)-stimulated maximal respiration were not altered in 4c9R in CA1(**Fig. 6I**). However, in DG basal OCR and FCCP-stimulated maximal OCR were significantly reduced in 4c9R mice compared to WT **(Fig. 6J).** Non-mitochondrial respiration after addition of rotenone and antimycin C was unchanged between genotypes in both CA1 and DG **(Fig. 6I, J)**. Our results thus indicate reduced basal and maximal mitochondrial respiration in 4c9R mice. Interestingly, when comparing CA1 to DG within a genotype, basal respiration was lower in DG compared to CA1 in 4c9R mice but no difference in basal respiration was observed between the two regions in wildtype mice (**Fig. 6K**). In addition, maximal respiration was significantly reduced in 4c9R mice compared to wildtype in DG (**Extended Data Fig. 4A)** but not in CA1 (**Extended Data Fig. 4B**). Spare/reserve capacity was not altered in DG or CA1 of 4cR mice (**Extended Data Fig. 4C, D**).

### Genomic fine mapping of behavioral phenotypes

Finally, we examined whether 3 or 4 copies of *Gldc* in the 3c9R and 4c9R mice are necessary and sufficient for the behavioral phenotypes previously seen in the 4c9LR mice which have 4 copies of the entire 9p24.1 segment (9LR). To this end, we performed behavioral assays pertaining to positive symptoms, negative symptoms and cognitive symptoms using mice with an increased copy number of *Gldc* only (3c9R, 4c9R) (see Methods and **Fig. 3A**), mice with an increased copy number of the other 9p24.1 genes except *Gldc* (3c9L, 4c9L) (see Methods and **Fig. 3A**), and the 4c9LR mice with four copies of all genes of the entire 9p24.1 segment as positive controls.

When testing latent inhibition (LI) to conditioned freezing, WT, 3c9L, and 4c9L mice pre- exposed to a tone before fear conditioning froze less than mice not pre-exposed to the tone, i.e., they displayed the LI effect. In contrast, in 4c9LR, 3c9R and 4c9R mice, the LI effect was absent **(Fig. 7A)**. As the LI effect is dependent on DG activity^41^, reduced glycine levels in the DG might contribute to the LI deficit. The experiments show that at least 3 copies of *Gldc* are necessary and sufficient for expression of the LI deficit, and an increase in the copy number of the other 9p24.1 genes does not affect LI.

**Figure 7.**
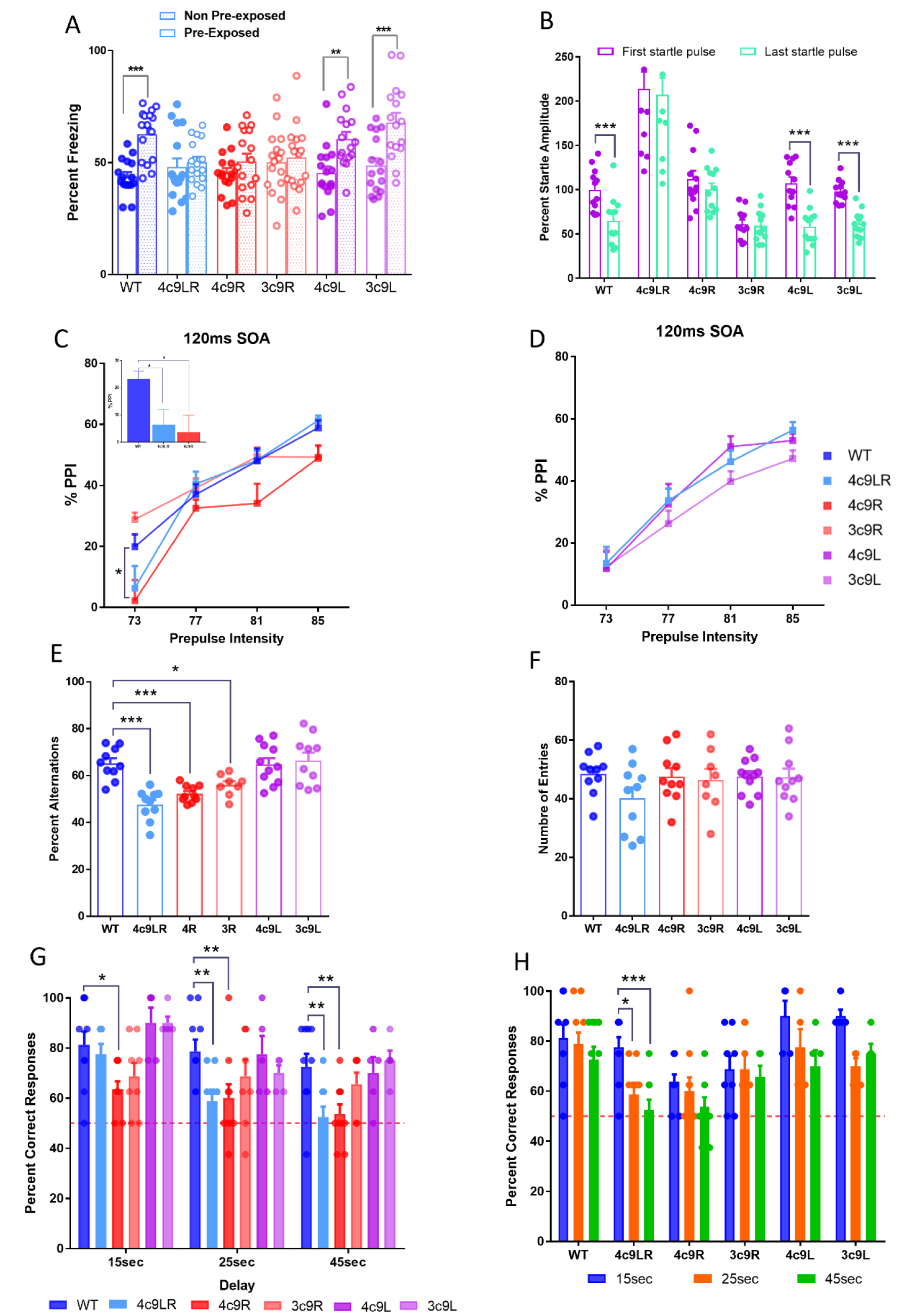
Effect of *Gldc* copy number variation on startle habituation, latent inhibition, prepulse inhibition and working memory. A. Latent inhibition to conditioned freezing. Bar graph comparing the percent freezing between two conditions, i.e., pre-exposed (white) and non-pre-exposed (matted) to a tone, within genotype for WT, 4c9RL, 4c9R, 3c9R, 4c9L and 3c9L mice (n=16 per bar (8 males, 8 females), One-way repeated measures ANOVA showed significant main effect of pre-exposure (F(11,165) =5.510, ***p<0.0001) indicating that mice not pre-exposed to a tone froze longer than mice pre-exposed to a tone. However, Bonferroni test showed significant difference in WT (t= 4.140, ***p<0.001), 4c9L (t= 3.338, **p<0.01) and 3c9L (t= 4.198, ***p<0.001), but not in 4c9LR, 4c9R and 3c9R mice. B. Habituation to acoustic startle. The startle responses for mice in response to the first and last startle pulses of each session are shown for each genotype (WT, 4cLR, 4cR, 3cR, 4cL, 3cL). Two way repeated measures ANOVA showed significant main effect of habituation process (F(1,66) =84.04, ***p<0.001) and significant main effect of genotype (F(5,66) =34.66, ***p<0.001) followed by main effect of interaction (F(5,66) =9.278, ***p<0.001). Bonferroni test revealed that there was significant habituation in wild type (t= 5.594, ***p<0.001), 4c9L (t=7.825, ***p<0.001) and 3c9L (t= 5.757, ***p<0.001) (n=10 with 5 male and 5 female mice). C, D. Line graph comparing prepulse inhibition to acoustic startle (PPI) at different prepulse intensities with SOA 120ms in WT, 4c9LR, 4c9R, 3c9R, 4c9L and 3c9L mice. At 120ms SOA, the two-way repeated measures ANOVA showed significant main effect of genotype (F(5,96) =3.441, **p<0.01), significant main effect of prepulse intensity (F(3,96) =112.7, ***p<0.001) on PPI and a significant interaction between prepulse intensity and genotype (F(15,96) =2.641, ***p<0.001). Further the Bonferroni test showed significant difference between WT vs 4c9R at 73dB prepulse intensity (C) (t= 3.028, *p<0.05) but not with the other genotype (D), (n=10 with 5 male and 5 female mice. The inset in C. shows the one-way ANOVA analysis for 73dB only at 120ms SAO, revealing a significant reduction of percent PPI in 4c9LR and 4c9R compared to wild type (One-way ANOVA followed by Bonferroni test, F(2,52)=4.270, *p<0.05; WT vs 4c9LR, t= 2.319, *p<0.05 and WT vs 4c9R, t= 2.708, *p<0.05). E, F. In the Y-maze spontaneous alternation test, the percentage of spontaneous alternations (E) (One-way ANOVA showed significant main effect of genotype on spontaneous alternations (F(5,53) =12.12, ***p<0.0001) and Bonferroni test showed significant difference between WT vs 4c9LR (t= 5.462, ***p<0.0001), WT vs 4c9R (t= 4.040, ***p<0.0001) and WT vs 3c9R (t= 2.702, *p<0.05), and the number of entries made into the arms (F) is shown for all genotypes (n=8-11, percent alternations). G, H, Water T-maze test. G. Bar graph showing percent correct choices made during test session comparing between genotypes at each delay duration mice (n=8-10). A the two-way repeated measures ANOVA showed significant main effect of delay durations (F(2,82) =18.78, ***p<0.0001) and significant main effect of genotype on correct choice (F(5,82) =5.126, ***p<0.0001). At 15 seconds delay, 4c9R mice had significantly reduced performance compared to WT mice (t= 2.810, *p<0.05). At 25- and 45-seconds delay, both 4c9LR (t=3.211, **p<0.01 and t= 3.211, **p<0.01) and 4c9R mice (t= 3.011, **p<0.01 and t= 3.011, **p<0.01) had significantly reduced performance compared to WT mice, respectively). The bar graph in H, based on the same data shown in G, shows percent correct choices made at different delay durations during test session comparing within each genotype (n=8-10). The two-way repeated measures ANOVA showed significant main effect of delay durations (F(2,126)=11.05, ***p<0.0001) and significant main effect of genotype (F(5,126) =9.348, ***p<0.0001). In 4c9LR mice, significantly reduced performance at 25 and 45 seconds compared to performance at 15 seconds (t= 3.011, *p<0.05 and t=4.014, ***p<0.001 respectively). In 4c9R and 3c9R mice, as the performance itself was low for all the delay durations, thus there was no difference between any delays. All data presented as mean ± SEM.

To assess the startle habituation effect, we compared startle response amplitudes to startle pulses alone presented at the beginning and at the end of a prepulse inhibition (PPI) session. We found that while wildtype, 3c9L an 4c9L mice displayed a reduction in the response to the last startle pulses compared to the first startle pulses, i.e., startle habituation, whereas 3c9R, 4c9R and 4c9LR mice responded indistinguishably to the first and last startle pulses, i.e., displayed a startle habituation deficit (**Fig. 7B)**. When assessing PPI to an acoustic startle stimulus, a measure of sensorimotor gating, PPI was reduced in 4c9LR (but not in 4c9R, 3c9L and 4c9L) mice at 73dB prepulse intensity with 60ms stimulus onset asynchrony (SOA) **(Extended Data Fig. 4G, H**). At 120ms SOA, at a 73 dB prepulse level 4c9R (but not 4c9LR, 3c9R, 3c9L and 4c9) mice displayed reduced PPI **(Fig. 7C, D**), indicating that 4 copies of *Gldc* lead to a PPI deficit. The response in the absence of a startle stimulus during the PPI session was not altered in any genotype compared to wildtype (**Extended Data Fig. 4I**). In summary, the results of startle experiments reveal that extra copies of the *Gldc* gene are necessary and sufficient for both PPI deficits and startle habituation deficits.

The Y maze spontaneous alternation paradigm was used to assess working memory. Mice with 3 or 4 copies of *Gldc*, i.e., 3c9R, 4c9R, and 4c9LR mice (but not 3c9L or 4c9L mice) displayed a reduction in the percentage of spontaneous alternations (**Fig. 7E)**, with the number of entries remaining unaltered between genotypes, suggesting no motivational alterations or differences in locomotor activity (**Fig. 7F**). In the non-matching to place water T-maze task, where mice were trained to alternate between two trials by rewarding correct alternations, in the testing phase intra-trial delays of 15 sec, 25 sec, and 45 sec were introduced that make the task more difficult. When compared to wildtype, 4c9LR mice and 4c9R mice displayed a reduction in the percentage of correct response rates. At 15 seconds delay, 4c9R mice had a significantly reduced performance compared to WT, and at 25- and 45-seconds delays, both 4c9LR and 4c9R mice had a significantly reduced performance compared to WT (**Fig. 7G**). To further clarify the effect of delays within genotype, we compared the within genotype performance between different delay durations (**Fig. 7H**). In 4c9LR mice, performance was significantly reduced at 25 and 45 seconds compared to 15 seconds, whereas in 4c9R and 3c9R mice, as the performance was low for all the delay durations including at 15 seconds delay, there was no difference between any delays. 3c9L and 4c9L mice performed indistinguishable from WT mice **(Fig. 7G, H)**. Our results show that 4 copies of *Gldc* are necessary and sufficient for disruption of working memory either in a spontaneous alternation task and in rewarded alternation task. The other 14 genes of the 9p24.1 CNV do not appear to contribute to this phenotype.

*S*chizophrenia frequently involves poor social functioning^42, 43^. We assessed whether mice with 9p24.1 copy number variation display differences in sociability and social novelty in the three-chamber social interaction test. In the sociability test, while WT, 3c9L and 4c9L mice spent more time with a stranger mouse under a cup (S1) rather than with an empty cup (E), the 4c9LR, 3c9R, and 4c9R mice spent almost equal time with S1 and E (**Fig. 8A, B**). In the subsequent social novelty test, while WT, 3c9L, and 4c9L mice spent more time exploring the novel stranger mouse (S2) compared to the now familiar mouse (S1), 4cLR and 4cR mice spent the same time with S1 and S2 mice, while 3c9R mice spent even significantly more time with S1 rather with S2 **(Fig. 8C, D**). The novelty preference index was reduced in 4c9LR, 3c9R, and 4c9R mice, compared to WT, but not in 3c9L and 4c9L mice (**Fig. 8E**). Additional behavioral characterization revealed no major genotypic differences in fear conditioning, forced swim test, open field test, and the novel object recognition test **(Extended Data Fig. 5A-H)**.

**Figure 8.**
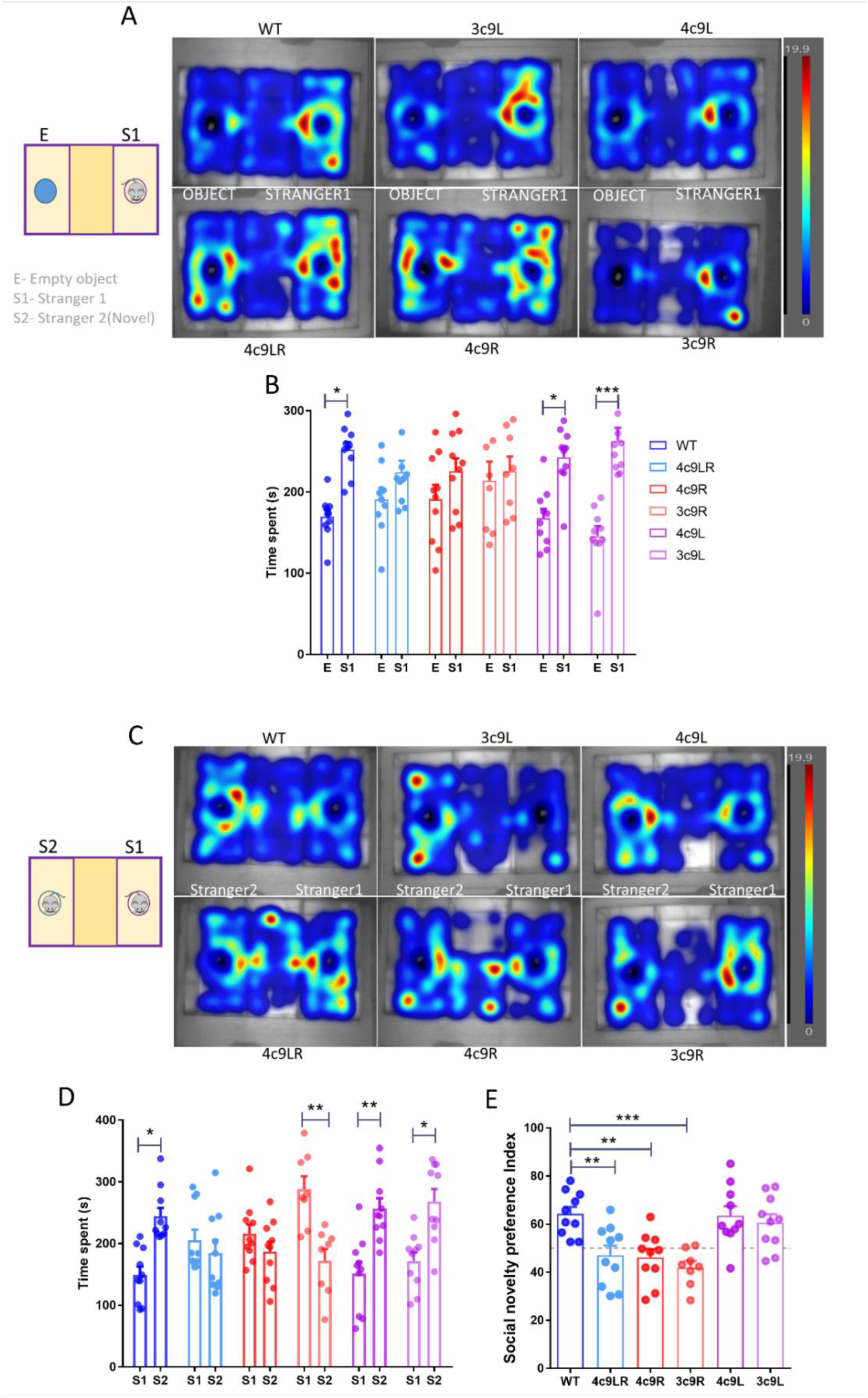
Effect of the GLDC copy number variation on social interaction behaviors. A, B. Sociability test. A. Heatmap showing the distribution of the average time spent by test mice for all genotypes during the sociability test. The schematic on the left side shows the three-chamber social interaction test setup for sociability assessment. B. Bar graph showing comparison of time spent with empty object (E) and time spent with stranger mouse-S1, an index of sociability (Two-way repeated measures ANOVA showed significant main effect of sociability, F(1,52) =27.27, ***p<0.0001), further Bonferroni test revealed significant differences between exploration of E vs S1 mice in WT (t= 3.030, *p<0.05), 4c9L (t= 2.765, *p<0.05) and in 3c9L (t= 4.318, ***p<0.001). C, D, E. Social novelty preference test. C. Heatmap showing the distribution of average time spent by test mice for all genotypes during the social novelty test. The schematic on the left side shows the representation of the three-chamber social interaction test setup for social novelty assessment. D. Bar graph showing comparison of time spent with stranger mouse-S1 (now familiar) and stranger mouse-S2 (Novel), an index of social novelty (Two-way repeated measures ANOVA showed a significant main effect of genotype on social novelty interactions, F(5,52) =3.08, *p<0.05) and a significant interaction between genotypes and social novelty interaction (F(5,52) =8.154, ***p<0.001). Further, Bonferroni test revealed significant interaction differences between S1 vs S2 mouse in WT (t= 3.142, *p<0.05), 3c9R (t= 3.422, **p<0.01), 4c9L (t= 3.480, **p<0.01) and in 3c9L (t= 3.195, *p<0.05). E. Bar graph showing the novelty preference index comparison between different genotypes (One-way ANOVA showed a significant difference in novelty preference, F(5,57) =7.947, ***p<0.001) between 4c9LR vs WT (t= 3.567, **p<0.01), 4c9R vs WT (t= 3.745, **p<0.01) and 3c9R vs WT (t= 4.350, ***p<0.001). n=8-10 (4-5 males and 4-5 females). All data presented as mean ± SEM.

## Discussion

In this study, we generated copy number variant mice mimicking a small supernumerary marker chromosome (establishing a 9p24.1 duplication/triplication CNV) found in patients with psychosis which showed phenotypic changes consistent with a schizophrenia-like pathophysiology. Functional genomic fine-mapping revealed that all observed phenotypes are due to increased copy number of *Gldc* but not an increased copy number of the other genes in the 9p24.1 CNV. These studies establish a potential link between a structural genomic variant found in patients, biochemical alterations such as increased GLDC expression and reduced BDNF levels, reduced extracellular glycine levels, a deficit in LTP, and working memory and other deficits (PPI, startle habituation, latent inhibition, and social interaction), which may be considered to constitute a schizophrenia-like biochemical and behavioral phenotype in mice. In patients with schizophrenia, BDNF is deceased in the postmortem brain^26^, and deficits in PPI^44, 45^, startle habituation ^46, 47^, latent inhibition^17, 18^ [a preclinical cognitive function paradigm of attention^48^], social functioning^42, 43^ and working memory^49^ have been reported. A reduced spine density is one of the most consistently observed neuropathological alterations in schizophrenia^33, 34^. Moreover, we found expression changes in genes or loci that have been linked to schizophrenia by GWAS, including miR-137^50^, which is involved in the regulation of synaptogenesis and synaptic plasticity^27, 51^, cortical morphology and cognition^36^, and ARL3^23, 38, 52^. All these phenotypes were recapitulated in the 4c9LR mice with the full 9p24.1 CNV as well as in the 4c9R mice with triplication of the *Gldc* gene alone. Furthermore, there was an enrichment of DEGs for genes that have been linked to ASD and NDD^20^. The increased expression of *Pyroxd2*, which is located in the mitochondrial inner membrane/matrix and interacts with complex IV subunit COX5B^21^, led to our finding that mitochondrial respiration is attenuated in the DG of 4c9R mice, in line with a report in organoids from patients with schizophrenia^40^. Our studies support the hypothesis that the small supernumerary marker chromosome found in the patients induces relevant neurobiological phenotypes in mice consistent with pathophysiological changes in neuropsychiatric disorders in humans and may thus be a contributing factor to the development of psychosis in these patients.

On a fundamental level, our study discovered that GLDC, which in the brain is expressed in astrocytes but not in neurons, modulates extracellular glycine levels in the DG but not in CA1, and that additional copies of *Gldc* reduce these glycine levels and thus attenuate LTP, which may underlie cognitive and/or other deficits.

## Online Methods

### Generation of mice with 9p.24.1 copy number variations

We have constructed mutant mice with a 0.9 Mb deletion or duplication on mouse chromosome 19 syntenic with the human 9p24.1 region by gene targeting and *trans*-allelic recombination *in vivo* **(Fig. 1A)**^53^. Briefly, we inserted two *lox*P sites into the mouse genome by gene targeting in V6.5 ES cells using insertion targeting vectors, one with homologous sequences from 19:29,302,809-29,306,803 (placing the “9L*lox*P” site at position 19:29,306,804 neighboring the *Rln1* gene) [in GRCm39], linearized at an internal *PflM1* site, the other with homologous sequences from 19: 30,189,922-30,196,026 (placing the “9R*lox*P” site at position 19:30,196,027, neighboring the *Gldc* gene), linearized at an internal *Xho1* site, and generated two separate mouse lines on a C57BL/6J background carrying these single *lox*P sites “flanking” the genomic segment that we wanted to delete and to duplicate. Correct targeting was confirmed by Southern blotting using external probes. Triple transgenic mice were then bred that carried the two *lox*P sites in *trans* and a *Hprt*-Cre transgene^54^. When breeding these mice to wildtype C57BL/6J mice, we obtained offspring that carried either the deletion (2.9%) or the duplication (2.9%), demonstrating *trans*-allelic recombination in the germline (for a schematic see **Fig. 1A**). Both deletion and duplication alleles (19:29,306,804-30,196,027, 889,223bp) were confirmed by diagnostic PCR assays and by comparative genomic hybridization **(Fig. 1B)**. Selectable neomycine resistance markers that were included in the targeting vectors have been removed using *Flp*-FRT recombination. The mouse nomenclature name for the mice carrying the 9p24.1 deletion is B6.Cg(129S1)-Del(19Rln1-Gldc)1Uru (deposited with MMRC, ID 68237), and the mouse nomenclature name for the mice carrying the 9p24.1 duplication is B6.Cg(129S1)-Dup(Rln1-Gldc)1Uru (deposited with MMRRC, ID 68238). The *lox*P site next to the *Rln1* gene was named “9L*lox*P” (L: left), the *lox*P site next to the *Gldc* gene “9R*lox*P” (R: right), and the genomic segment between these two *lox*P sites “9LR”. 9LR mice were backcrossed to C57BL/6J for at least 5 generations.

We then inserted a *lox*P site between the *Uhrf2* and *Gldc* genes using CRISPR/Cas-9. The oligonucleotide from which the guide RNA was derived had the sequence CTGACATTAATTGAGACCGTGGG (19:30,072,176-30,072,154(-)). The resulting construct was electroporated into zygotes from the C57BL/6J background. Embryos were cultured overnight and transferred to pseudopregnant females (University of Pittsburgh). The presence of the *lox*P site was confirmed by PCR. This *lox*P site was named “9I” (I: intermediate). We then bred mice carrying the “9L*lox*P” *lox*P site, the “9I*lox*P” *lox*P site and the Hprt-Cre transgene and bred these mice with C57BL/6J wildtype mice and – via *trans*-allelic recombination – obtained offspring that carried a duplication of the segment between these *lox*P sites, named “9L”. In a similar way, we bred mice carrying the “9I*lox*P” *lox*P site, the “9R*lox*P” *lox*P site and the Hprt-Cre transgene and bred these mice with C57BL/6J wildtype mice and obtained offspring that carried a duplication of the segment between these loxP sites, named “9R”. The “9L” and “9R” genotypes were identified by PCR. The Hprt-cre transgene was bred out. Experiments were performed with the following three genotypes: “9LR” mice containing a duplication of the entire *Rln1*-*Gldc* segment, “9R” = B6.Cg(129S1)-Dup(Gldc)1Uru mice containing a duplication of just the *Gldc* gene, and “9L” = B6.Cg(129S1)-Dup(Rln1-Uhrf2)1Uru mice containing a duplication of the *Rln1*-*Uhrf*2 segment, i.e., of the *Rln1*-*Gldc* segment excluding the *Gldc* gene.”3c9R” (“3c9L”) harbor the duplication on one chromosome 19 (also called “duplication”) and thus have 3 copies of this genomic segment, “4c9LR”, “4c9L”, and “4c9R” mice harbor the duplication on both chromosome 19 (also called “triplication”) and thus have 4 copies of this genomic segment **(Fig. 3A)**. Mice are on the C57BL/6J background (at least 8 backcrosses). ARRIVE guidelines were followed.

### Subjects

Mice with copy number variants in the (human) 9p24.1 region were maintained on a C57BL/6J background at McLean Hospital and at the University of Illinois. Mice approximately 2-6 months of age were used in experiments. Wildtype mice were derived from mutant crosses. Mice were typically genotyped by copy number PCR from tail biopsy samples prepared using a Puregene Core Kit A (Cat no-158267, Qiagen, MD). Copy number PCRs were performed using commercial kits designed to detect the Gldc and Uhrf2 genes (Mm00496811_cn GLDC (#4400291), Mm00735356_cn Uhrf2 (#4400291) and TaqMan™ Copy Number Reference Assay, mouse, Tfrc (#4458366) Thermo Fisher Scientific, MA). The mice were typically housed in groups of four to five per cage in standard conditions (23 ± 1 °C, 50 ± 5% humidity) with 12-h light-dark cycle (lights on from 07.00-19.00). Food pellets and water were provided *ad libitum*. Mice were mostly tested during the light period of the light/dark cycle, except for the light/dark choice, novelty-suppressed feeding and elevated plus maze tests, when mice were testedreac during the dark period. All experimental procedures were conducted in accordance with the Guide for the Care and Use of Laboratory Animals (National Research Council) and approved by the IACUCs of McLean Hospital, the University of Illinois, the University of Pittsburgh and the University of Bonn. For the experiments performed at McLean Hospital described in **Fig. 1** and **Extended Data Fig. 1**, male mice were used. For all other tests, both male and female mice were used. As no differences were detected between males and females (with the exception of **Extended Data Fig. 7F**), they were collapsed into one group.

### Prepulse inhibition and startle habituation

Prepulse inhibition (PPI) was determined using the SM100 Startle Monitor System (Kinder Scientific, USA), consisting of a sound-attenuated startle chamber and Startle Monitor software. Animals were placed in an adjustable Plexiglas holder, allowing limited movement but not providing restraint, positioned directly above a sensing platform. Each animal underwent 15 min of habituation with the holder for two days in the home cage, followed by habituation to the testing chamber with white noise for 5 min on the next day. On test day, each test session consisted of 5 min acclimatization with background white noise (70 dB), followed by presentation with 6 startle pulses of 120 dB [intertrial interval (ITI): 20 s]. Animals were then subjected to five types of trials presented 12 times each in a pseudorandom manner: pulse alone, prepulse + pulse, or prepulse alone (prepulse 3, 6, 9 or 12 dB above background). ITI varied from 10 to 20 s (average 15), two stimulus onset asynchrony (SOA) intervals were 60ms and 120 ms, prepulse length 20 ms, and pulse length 40 ms. Each PPI session ended with 6 startle pulse presentation of 120 dB. The PPI test lasted about 35 min.

PPI was calculated as % PPI for each prepulse intensity as: [(pulse_alone - prepulse + pulse/pulse_alone) × 100]; with a lower percentage indicating decreased PPI. The startle magnitude was calculated from the average of pulse alone trials. Habituation was assessed by comparing the averages of first (6) and last (6) sets of startle pulse alone trials.

### Three-Chamber Social Interaction Test

Sociability and social novelty were evaluated in a three-chamber social interaction test. The apparatus is a rectangular three-chamber box in which each chamber is 20 x 45 cm with a height of 40 cm. Walls and dividing walls are made from clear Plexiglas, with a sliding door middle section with 5 cm width and 5 cm height which allows free access to each chamber. The subject mice were habituated to the central compartment with closed doors for 5 min. After the habituation phase, subjects were tested in the sociability task that consisted of exploring either a conspecific stranger mouse 1 (Sex-matched one-month younger mouse inside an inverted cup) or an empty inverted cup located in the opposite compartment with an exploration time of 10 min. After this session and a 5 min break, the social novelty task was performed for 10 min. The empty inverted cup is replaced with an inverted cup holding a novel stranger mouse 2. Mice under the cups are sex-matched to and approximately one month younger than the experimental mice. The apparatus is cleaned with 70% ethanol after each trial. The time spent in each compartment was measured. The data were recorded and analyzed using the EthoVision XT15 (Noldus Inc., Leesburg, VA, USA).

### Latent Inhibition to conditioned freezing

We used a modified version of the protocol by E Engin et al^41^. Mice were assigned randomly to a “preexposure” or “non-preexposure” group on day 1. The mice in the preexposure group were placed in Context-A (floor insert, white chamber light, 0.5% benzaldehyde as olfactory cue) and were presented with 30 tones (20 s, 90 dB, 5000 Hz) with 30 s intervals. The non-preexposure group were placed in the same chamber but no tone was presented. On day 2, all mice were placed in Context B (no inserts, small stimulus light on wall, 1% acetic acid as olfactory cue) with grid floors and were subjected to five tone (20 s, 90 dB, 5000 Hz)–shock (2 s, 0.7 mA) pairings with 60 s intervals. On day 3, all mice were placed in Context C (floor and ceiling inserts, white chamber light, 1% peppermint extract as olfactory cue), the tone was presented after 3 min for 6 min and freezing behavior was recorded and analyzed using the video freeze software (Med-Associates Inc., Fairfax, VT).

### Y-maze spontaneous alternation

Working memory was tested using a symmetrical Y-maze with three arms (30 cm x 8 cm x 15 cm) at 120° angles to each other. Briefly, the mouse was placed into the end of one arm and allowed to explore freely through the maze during an 8-min session. The sequence (pattern) and total number of arms entered were recorded. An arm entry is considered to be complete when the hind paws of the mouse had been completely placed in the arm. The percentage of alternations is the number of alternations divided by the total number of arms entered multiplied by 100. As the re-entry into the same arm was not counted for this analysis, the chance performance level in this task was 50% in the choice between the arm mice visited more recently (non-alternation) and the other arm the mice visited less recently (alternation).

### Paired-trial Variable-delay Water T-maze task

Working memory was tested by measuring acquisition in a nonmatching-to-place test. The water-T-maze used was constructed from gray plexiglas, with the following dimensions: length of stem 70 cm, length of arms 30 cm, channel width of 11 cm, and wall height of 20 cm. Water (maintained at 23°C±1°C) was added to the apparatus up to a height at which a submerged platform (5 x 10 x 9 cm) would be concealed 1 cm below the surface at the end of the arms. Data were collected manually by a single observer. Two days of habituation to the maze were followed by 8-12 training days, and the last two days are test days.

At the beginning, mice were given 2 day of habituation/training consisting of 4 sets of forced alternation between the left (L) and right (R) arms by switching a removable barrier between the arms. In this way, mice were habituated to the procedure and learned to reliably swim down the T-maze to find the platform at the end of the specific open arm. During the forced alternation training sessions, 100% of mice from all genotypes successfully reached the platform with none showing deficits in swimming ability.

### Training

The next day, an additional habituation session (2 sets of forced swimming to left (L) and right arm (R)) was performed followed by a series of 14 test trials. Each trial consisted of two runs. The first run involved a forced choice, with one arm blocked and the other containing the platform. Once the mouse reached the platform, it was left there for 5 seconds and then picked up and returned to the starting point to begin the second run, the free choice phase of the trial. During this phase, the barrier was removed, and the platform was moved to the opposite arm. In this run, if the mouse swam to the arm opposite to the initial one, i.e. the arm now containing the platform, it was considered a success and it was rewarded by being allowed to stay on the platform for 10 sec. If the mouse swam to the arm without a platform, i.e. not alternating, it was considered a failure and, as punishment, the mouse was confined and forced to swim in that arm for 15 sec. Mice were not allowed to reverse course, and were deemed to have selected an arm when the hind feet crossed into it. Mice were then given a 2-minute intertrial interval. Left-Right order for 14 trials per day was assigned randomly to each mouse. This procedure was repeated daily until the mouse reached 70% correct choice for three consecutive days or up to the next 10 days to conclude the training. The number of correct choices by total trials indicates the learning ability. During the forced alternation training sessions, 100% of mice from all genotypes successfully reached the goal platform with none showing deficits in swimming ability.

### Testing

there are two additional habituation rounds with choice and punishment, one on each side. Then, there are 12 test rounds, 4 rounds per intra-trial delay. The forced trials are as normal, and then there is either a 15, 25 or 45 second intra-trial delay. The longer the intra-trial delay, the more difficult the task is for the mouse. Percent correct choice is calculated separately for the testing trials. Data were analyzed using a two-way repeated measures ANOVA and the Bonferroni test.

### Novel object recognition (NOR) test

The NOR test was performed to assess recognition memory. The experimental apparatus consisted of a black rectangular open field (40 × 40 × 30 cm) apparatus. On the first two days of habituation each mouse was allowed to habituate to the empty chamber for 10 min. On the test day, during the training phase mice were placed in the apparatus in the presence of two identical objects and were allowed to explore for 10 min. After an hour in their home cage, mice were placed again in the apparatus for 10 min, where one of the objects had been replaced with a novel object. The objects consisted of two small plastic boxes with different shapes, both approximately of the same height. In both tasks, the arena was cleaned with 70% ethanol to eliminate previous mouse smells. During object recognition testing, the interaction time with the familiar object and the novel object was recorded by the EthoVision XT15 video-tracking system using the multiple body point module (Noldus Inc., Leesburg, VA, USA) and analyzed. A discrimination index was calculated for each mouse as follows: time spent exploring the novel object / total time spent exploring both objects *100. Values for the discrimination index ranged from 0 to 100, where a score closer to 100 indicated greater preference for the novel object, while a score closer to 0 indicated preference for the familiar object.

### Forced Swim Test (FST)

The FST apparatus was a transparent Plexiglas cylinder (20 cm diameter, 30 cm height). The cylinders were filled with 25 ± 1°C water to a 20 cm depth to prevent animals’ tails from touching the bottom of the tank. The animals were placed in the cylinders for 6 min (Extended Data Fig. 1) or 10 min (Extended Data Fig. 5). Activity was recorded by a video camera placed directly facing the side wall of the cylinder for later scoring. Animals were dried with paper towels after each exposure and returned to preheated home cages. Duration of immobility, i.e., when the mice perform the minimum movement necessary to stay afloat, was measured. Scoring was performed manually in bins with each bin of 5 seconds of continuous floating.

### Fear Conditioning

Mice were subjected to a trace fear conditioning protocol adapted from E Engin et al^41^ with slight modifications. On the first day of the experiment, the mice were placed in a conditioning box (Med-Associates Inc., Fairfax, VT) with grid floors, and subjected to 5 tone (20 s, 90 dB) – shock (2 s, 0.5 mA in Extended Data Fig. 1; 2 s, 0.7 mA in Extended Data Fig. 5) pairs with 20 s trace period between the tone and the shock (trace fear conditioning) or the shock coinciding with the end of the tone (delay fear conditioning). 24 hr later, they were returned to the same context for 180 s and freezing was recorded (contextual conditioning). On day 3, the mice were placed in a different context, the tone was presented for 6 min and freezing behavior was recorded and analyzed using the Video Freeze Software (Med-Associates Inc.).

### Elevated plus maze

The elevated plus maze had two open arms (35 x 6 cm) and two enclosed arms (35 x 6 x 20 cm) separated by a center area (5 x 5 cm) at an elevation of 1 m above the ground. Light intensity was approximately 30 lux in the open arms. Mice were inserted in the center facing one of the open arms for 5 min testing sessions, which were analyzed with Noldus Ethovision XT tracking software. The percentage of open time in the open arms was calculated as open arm time/5min x 100.

### Light/dark choice text

The apparatus for the light/dark choice test consisted of a lit (200 lux) chamber (28 x 28 x 31 cm) and a dark (<10 lux) chamber (14 x 14 x 31 cm) connected by a small opening. Mice were placed into the dark chamber and allowed to explore the apparatus for 6 min. Sessions were recorded and analyzed for time spent in light with the Noldus Ethovision XT software.

### Novelty-suppressed feeding

Food-deprived (36 h) animals were inserted in a corner of a clear plexiglas box (42cm x 42 cm x 31 cm) at 100 lux illumination, with clean bedding and an inverted petri dish with a food pellet placed in the center. The latency to bite the pellet and to sit and eat the pellet were recorded. Sessions were for 3 min, with animals being removed after they began eating.

### Sucrose Preference test

Mice were allowed to choose between 0.5% (w/v) sucrose and water over four days. The position of the bottles was exchanged after two days. The preference to sucrose was calculated by total sucrose consumed divided by total liquid consumed multiplied by 100.

### Cellular fractionation protocol

Adult (2-4 month-old) male and female mice were sacrificed by cervical dislocation and their brains were quickly removed. Immediately the brain was rinsed in cold PBS, and was coronally sectioned into two halves placing blade at approximately 2.0 mm posterior to bregma (Paxinos and Franklin, 2001). Initially PFC was dissected from the anterior half and then the posterior portion was used to dissect and collect both the lobes of the hippocampus. The tissue was flash frozen in dry ice and stored at −80 °C until analysis.

The fractionation procedure used here was modified from a previously published protocol ^56–58^. Tissue was homogenized with powergen-125 instrument in 0.4ml of ice-cold RIPA lysis buffer (150 mM NaCl, 1% Nonidet P-40, 0.5% Sodium Deoxycholate, 0.1% SDS, 50mM Tris (pH 7.4)) containing 1mM HEPES (pH 7.4) with freshly added protease and phosphatase inhibitor cocktail- AEBSF, Aprotinin, Bestatin hydrochloride, E-64, Leupeptin, Pepstatin A, Cantharidin, (–)-p-Bromolevamisole oxalate, Calyculin A (PPC1010-5ml, Sigma). The homogenized tissue was centrifuged at 1,000×g for 10 minutes at 4 °C to separate a pellet enriched in nuclear components (P1). The supernatant was then centrifuged at 15,000×g for 20 minutes at 4 °C to obtain a clarified cytosolic fraction (CF) and a pellet enriched with synaptosome membranes. The pellet (P2) was resuspended in ice-cold RIPA buffer containing 150 mM NaCl, 1% Nonidet P-40, 0.5% Sodium Deoxycholate, 0.1% SDS, 50mM Tris (pH 7.4), and 20mM HEPES, then sonicated on ice for three times with a 60 second interval, each time with 6 strokes (samples were placed on ice during and between sonication), resulting in a crude synaptosome fraction which was used for Western blots. The protein was immediately stored at −80 °C until analysis. Protein estimation was performed with Pierce™ BCA Protein Assay Kit (Thermo Scientific) and samples were prepared for western blotting.

### Mitochondrial respiration measurements

Mice were euthanized by decapitation and brains were rapidly removed (within 60 s of decapitation) and immersed in ice-cold (4–5°C) artificial cerebrospinal fluid (aCSF; 120 mM NaCl, 3.5 mM KCl, 1.3 mM CaCl2, 1 mM MgCl2, 0.4 mM KH2PO4, 5 mM HEPES, and 10 mM D-glucose; pH 7.4) that had been oxygenated for 1 h using carbogen (95% O2: 5% CO2). Coronal Sections (200 μm) were prepared using a vibratome 1000 plus (IMEB Inc, Sa Marcos, CA) with the oxygenated chilled aCSF on stage and blade, then transferred to a holding chamber containing continuously oxygenated aCSF on ice (∼ 4 °C).

These coronal sections were individually transferred to a biopsy chamber containing fresh oxygenated aCSF. Stainless steel WellTech Rapid-Core biopsy sampling punch needles (500 μm diameter; TED PELLA Inc., Redding, CA) were used to excise the hippocampal sub-region tissue punches. Hippocampal punches were collected from both hemispheres for each anatomical location using consecutive coronal sections.

Punches were ejected directly into an XFe96 Cell Culture Microplate (101085-004; Agilent Technologies, Santa Rosa, CA) based on a pre-determined plate layout. Each well contained 180 μl room temperature assay media (aCSF supplemented with 0.5 mM pyruvate). After loading all biopsy samples, each well was visually inspected to ensure that the punches were submerged to the bottom of the well. The XFe96 Cell Culture Microplate was then incubated at 37 °C for approximately 30 min.

During this incubation period, 10 X concentration of assay drugs (prepared in aCSF) were loaded into their respective injection ports of a hydrated (overnight in distilled water) Seahorse XFe96 Extracellular Flux Assay sensor cartridge. The sensor cartridge containing the study drugs was then inserted into the analyzer for calibration. Once the analyzer was calibrated, the calibration plate was replaced by the microplate containing the tissue punches and the assay protocol initiated.

The tissue respiration assay protocol consisted of a minimum of three cycles of oxygen consumption rate (OCR) measurements for each measurement period. Each cycle consisted of a 2 min ‘mix’ period and a 2 min ‘wait’ period, followed by a 3 min ‘measure’ period. Three cycles were used to obtain a basal OCR, 3 cycles were used to assess the effect of the F1-Fo ATP synthase inhibitor oligomycin, 3 cycles were used to evaluate the effect of the uncoupler FCCP (Carbonyl cyanide-4 (trifluoromethoxy) phenylhydrazone), and 3 cycles were used to measure mitochondria-associated respiration following injection of antimycin A / rotenone. For optimization of tissue punch diameter and drug concentrations the same procedures were applied.

Upon completion of the assay, the OCR curve generated by each well was examined using Agilent’s Wave 2.6.0 software. The samples that failed to respond to FCCP/pyruvate were excluded, as this represents a failure of the assay, possibly due to movement of the tissue punch during mixing cycles. In a typical experiment, 5% or fewer wells were excluded. Experiments for mitochondrial function analysis were replicated in three wells per tissue type per mouse. The data are imported into Excel format and results were averaged for each genotype. We calculated the OCR for all inhibitors sequentially added, separately for basal mitochondrial respiration (OCR in the absence of any inhibitors), maximal respiration (basal respiration + spare capacity), and the spare (or reserve) capacity (FCCP coupled OCR – basal OCR).

### Golgi Staining and Dendritic Spine Analysis

Extracted brains were subjected to Golgi–Cox impregnation and staining of neurons as per the manufacturer’s instructions (SuperGolgi kit, Bioenno Tech). Whole brains were impregnated in solution-A at room temperature for 12 days. After immersion in the post-impregnation solution-B for 2 days, brains were serially sectioned at 150 μm using a vibratome and the sections were mounted on 0.1% gelatin-coated slides, washed in 0.01 M PBS buffer (pH 7.4) with Triton X-100 (0.1%), and stained solution-C. After staining and washing, slides were gradually dehydrated through a series of increasing alcohol concentrations, then with xylene and finally mounted in Eukitt ® Quick-hardening mounting medium (Sigma Aldrich) for analysis.

Spine analyses were conducted in the dentate gyrus region of the hippocampus. For spine quantification the untruncated secondary dendrites were selected with dendritic spines visually distinct from one another having clearly defined spine heads. One dendritic segment of at least 10 um in length was analyzed per neuron, and 2-3 neurons were analyzed per brain. All images were captured with the NeuroLucida system at 100× magnification in Zeiss Axio Imager A1 inverted microscope.

To quantify different type of spines, the geometry of the spine shapes was examined to classify them with an unbiased method. In brief, measurements of the head width and neck length and the length–width ratio (LWR) were performed to determine spine types according to the following criteria: filopodia (length > 2 µm), long/thin (length > 1 µm), thin (LWR > 1 µm), stubby (LWR ≤ 1 µm), and mushroom (width > 0.6 µm) ^59^. Analysis of density and morphology of dendritic spines was performed by using the Reconstruct software (https://synapseweb.clm.utexas.edu/software-0).

### Determination of the glycine decarboxylase (GLDC) enzyme activity

GLDC is the first and rate limiting enzyme of glycine cleavage system (GCS) also called glycine decarboxylase complex. We used Glycine Cleavage System (GCS) Assay Kit (Cat #: E-137, BRS, University at Buffalo, NY) which is based on the reduction of the tetrazolium salt INT in a NADH-coupled reaction to formazan, that exhibits an absorption maximum at 492 nm allowing for sensitive detection of GLDC activity via GCS complex activity.

### Estimation of glycine levels in acute hippocampal slices using ratiometric optical FRET sensor GlyFS

Experiments were performed as previously described^29, 60^ with minor modifications. Animals were housed under 12 h light/dark conditions and were allowed ad libitum access to food and water. Acute hippocampal slices were prepared from 2–3-month-old wild type and 4-copy *Gldc* male and female mice in full compliance with national and institutional regulations (Landesamt für Natur, Umwelt und Verbraucherschutz Nordrhein-Westfalen and University of Bonn Medical School) and guidelines of the European Union on animal experimentation. All efforts to minimize number of animals used and their suffering were made.

Horizontal slices (300 µm) containing hippocampal formation were prepared in an ice-cold slicing solution containing (in mM): NaCl 60, sucrose 105, KCl 2.5, MgCl2 7, NaH2PO4 1.25, ascorbic acid 1.3, sodium pyruvate 3, NaHCO3 26, CaCl2 0.5, and glucose 10 (osmolarity 300–310 mOsm) and kept in the slicing solution at 34 °C for 15 min before being stored at room temperature in an extracellular solution containing (in mM) NaCl 131, KCl 2.5, MgSO4 1.3, NaH2PO4 1.25, NaHCO3 21, CaCl2 2, and glucose 10 (osmolarity 297–303 mOsm, pH adjusted to 7.4). This solution was also used for recordings.

For anchoring of GlyFS in brain tissue, cell surfaces within acute slices were biotinylated using a previously published procedure^29, 60^. Briefly, the slice storage solution was supplemented with 50 µM Sulfo-NHS EZ Link Biotin (Thermo Fisher) for 45 min before washing and further storage. For experiments, slices were transferred to a submersion-type recording chamber and superfused with extracellular solution at 34 °C. All solutions were continuously bubbled with 95% O2/ 5% CO2.

GlyFS (final concentration 100 µM) was dissolved in PBS (pH 7.4) and 12.5 µM streptavidin (Thermo Scientific) was added. GlyFS was pressure injected in stratum radiatum (SR) of CA1 or in the inner one-third of molecular layer of the DG via micropipette (2–4 MΩ when filled with the sensor solution) inserted ∼70 µm deep into the tissue under visual control. The same pipette was used for fEPSP recordings. GlyFS-injected slices were allowed to recover for 10–15 min before recordings.

GlyFS fluorescence was visualized by two-photon excitation (2PE) fluorescence microscopy. We used a FV10MP imaging system (Olympus) or Scientifica 2PE fluorescence microscope (Scientifica) optically linked to a femtosecond pulse laser (Vision S, Coherent, λ = 800 nm) and equipped with a 25× (NA 1.05) or 40× (NA 0.8) objectives (Olympus). The laser power was adjusted for depth in the tissue to obtain, on average, a fluorescence intensity corresponding to that of 2-3 mW laser power at the slice surface. Fluorescence of ECFP and Venus fluorescent protein was separated using appropriate band pass filters and dichroic mirrors and detected with photomultiplier tubes (PMT). PMTs gains were constant throughout the whole set of experiments. GlyFS fluorescence was acquired either in frame scanning mode (512x512, 3 frames per imaging stack) for estimation of resting levels of glycine or in line scan mode (0.2-0.3 kHz) in experiments with high frequency stimulation (HFS, see below). Electrophysiological data were recorded using MultiClamp 700B amplifiers, digitized (20 kHz) and stored for offline analysis. For stimulation experiments, a bipolar concentric stimulation electrode was placed in the CA1 or in the DG in the same layer as the recording microelectrode to stimulate Schaffer collaterals (SC) or medial perforant path (mPP) respectively. The stimulus intensity was adjusted to evoke half-maximal fEPSPs. HFS was performed at a frequency of 100 Hz for 1 s. Line scans were time synchronized with electrophysiological recordings and were acquired 1 s immediately before the HFS, during the 1 s HFS and 1 s immediately after the HFS.

Image analysis was performed in FIJI/ImageJ (NIH) and custom written Matlab scripts (Matlab R2020b, Mathworks). Images within one image stack were averaged. After background correction, the ratio of ECFP and Venus fluorescence was calculated from the fluorescence intensities in the respective channels. Background intensity was subtracted for each channel.

To estimate the glycine levels in slices, ECFP/Venus fluorescence intensity ratios R0 and RMAX were calculated, where R0 and RMAX represent the resting and maximal ratios for a glycine concentration of 5 mM. Exogenous glycine was applied via the bath perfusion (5 mM).

Statistical analyses were performed in Excel (Microsoft), Origin Pro (OriginLab Corporation) and MATLAB (Mathworks). Data are presented as mean ± s.e.m. Comparisons of populations were tested using Student’s t-tests or non-parametric tests if data sample showed non-Gaussian distribution as indicated. Data samples were tested for normality using the Shapiro-Wilk test.

### Electrophysiology

Hippocampal slices (400 µm in thickness) were prepared from genetically engineered mice with two copies (wild type, WT) or four copies (4c9R) of the glycine decarboxylase gene with vibratome in cold oxygenated cutting solution containing (in mM) 252 sucrose, 2.5 KCl, 1.0 CaCl2, 5.0 MgSO4, 1.25 NaH2PO4, 26.0 NaHCO3, and 10 glucose. Slices were kept in the artificial cerebrospinal fluid (ACSF) containing (in mM) 119 NaCl, 2.5 KCl, 2.5 CaCl2, 1.0 MgSO4, 1.25 NaH2PO4, 26.0 NaHCO3, 10 Glucose, and equilibrated with 95% O2 and 5%CO2 (pH 7.3–7.4) at room temperature for one hour before recordings. The recordings were conducted in the presence of bicuculline (10-20 µM, GABAA receptor antagonist) in perfusion medium at 22-24°C. Synaptic transmission and plasticity were assessed at synapses in the medial perforant path to the dental gyrus (mPP-DG) or the Schaffer collaterals to the CA1 (CA3-CA1) field of the hippocampus, respectively. Field excitatory postsynaptic potentials (fEPSPs) were recorded in the middle molecular layer of the dentate gyrus or the stratum radiatum of the CA1 area with a glass pipette (3-4 MΩ in resistance) filled with the extracellular solution. Synaptic responses were evoked by stimulation of the medial perforant path fibers (mPP)^62^ or Schaffer collateral fibers with a concentric stimulation electrode ^63, 64^. Paired pulse ratio was calculated as the ratio of the rising slope of the second fEPSPs to the first fEPSPs at the interstimulus intervals of 50, 150 and 300 ms. LTP was induced by a 1-s stimulation train at 100 Hz after recording basal synaptic transmission for 20 minutes at the stimulation frequency of once every 30 s. The recordings were continued for an additional 50 minutes after the induction of LTP. The magnitude of LTP was estimated 40 min after the induction.

### Western Blot

Western blot analyses were performed on cytosolic fraction (CF) (n=5 per genotype, 3 males + 2 females) and synaptoneurosomal fraction (SF) (n=5 per genotype, 3 males + 2 females) protein fractions. Prior to gel loading, CF and SF samples were heated to 95 °C for 5 minutes, 20 ug of CF and SF protein samples (along with protein ladder in first well) were electrophoretically separated on an SDS 10-11% polyacrylamide gel and transferred to a precisely cut PVDF membranes. PVDF membranes (Immobilin-P, Millipore Inc., St. Louis, MO, USA) were blocked with 5% skimmed milk powder (LP0031, OXOID, KS, USA) in 0.05% Tween-20/Tris-buffered saline and then incubated with primary antibody overnight in cold room (∼4^0^ C). After incubation with goat anti-rabbit (1:5000; Cell Signaling) or rabbit anti-mouse (1:3000; Cell Signaling) horseradish peroxidase-conjugated secondary antibodies, for 1 hour and washed with TBS. Semi-quantitative assessment of protein bands was performed by developing the blots for 2 min with chemiluminescence substrate using SuperSignal™ West Femto Maximum Sensitivity Substrate (Thermo Fisher Scientific, Waltham, MA, USA) and imaged the blots that was executed by Protein Simple Fluor Chem R (Protein Simple, USA. Densitometric quantification was performed using Image-J software. Each blot was used to detect multiple proteins. **The list of primary antibodies as below**.

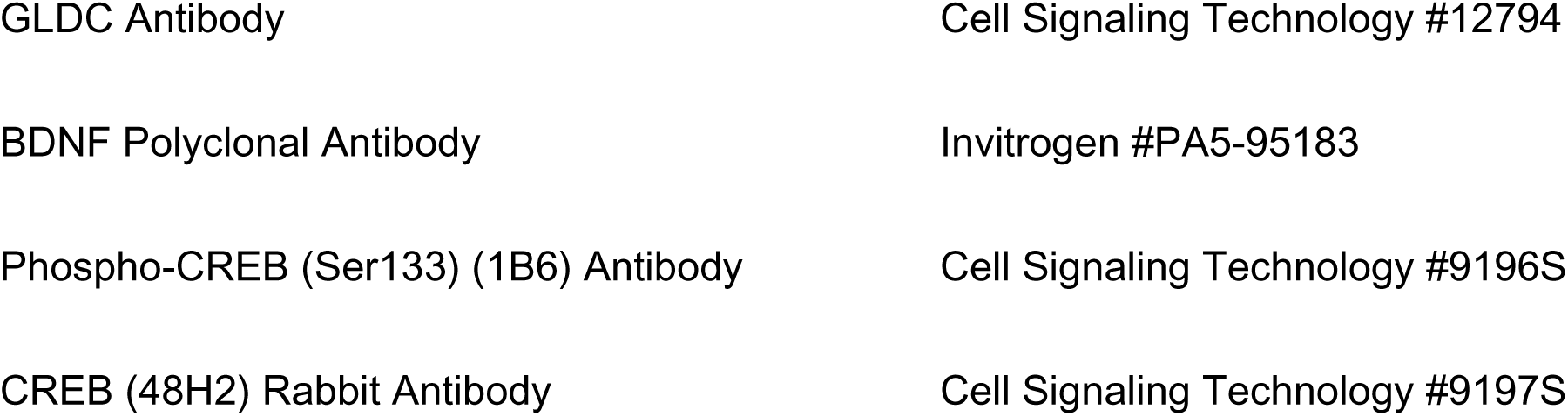
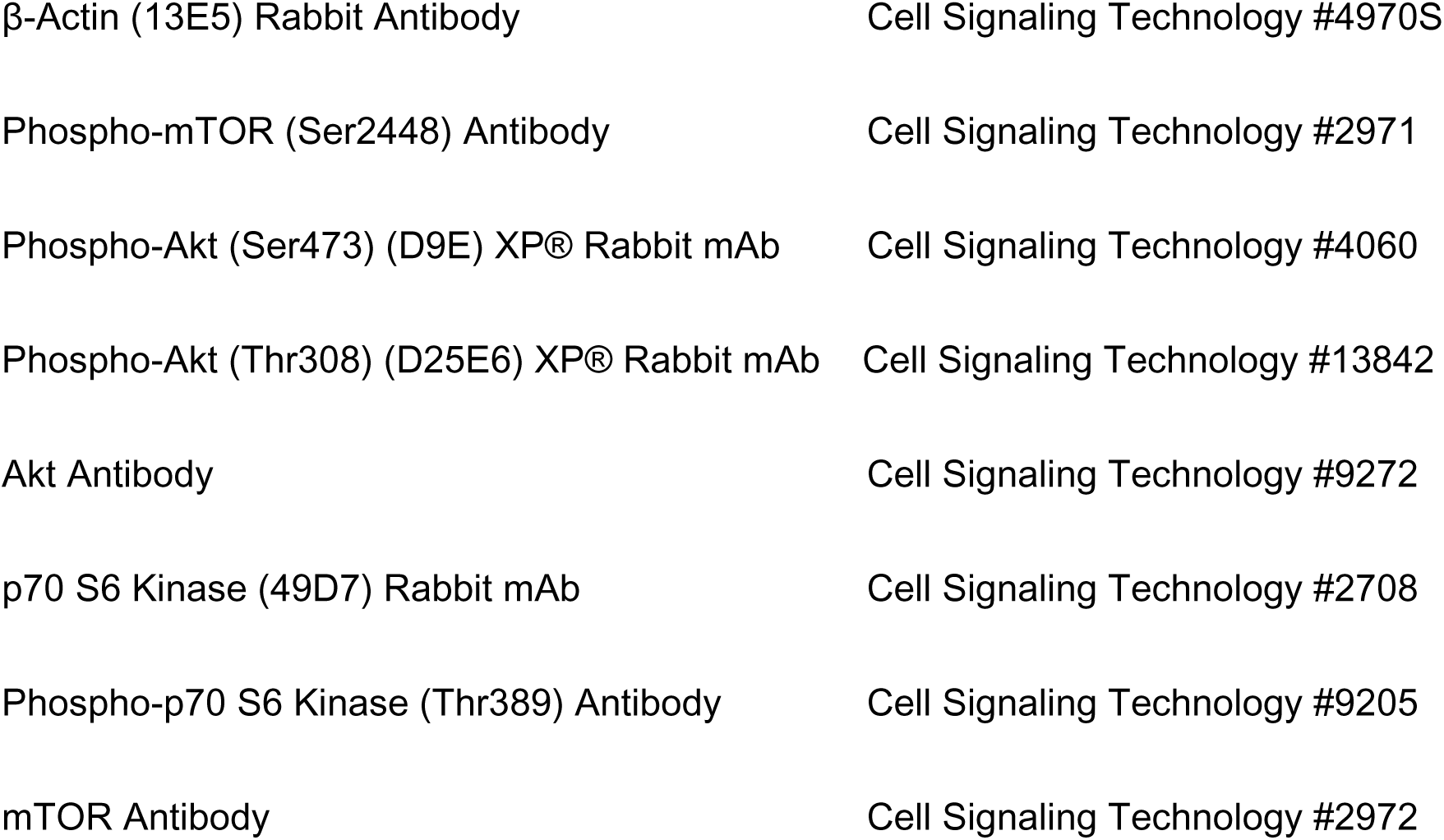

### Quantitative real-time reverse transcription polymerase chain reaction (qRT-PCR)

RNA was isolated from the hippocampus of WT and 4c9R mice (n = 5/genotype) using the miRvana miRNA isolation kit (Ambion, Austin, TX, USA). cDNA for each RNA sample (5-μg input) was generated using the High-Capacity cDNA Reverse Transcription kit (Applied Biosystems). For the microRNAs (miR-132, miR-137 and sno-202), quantification was performed with Qubit assay (Q32880) using the Qubit 4.0 Fluorometer, then cDNA (100-ng input) was prepared using the TaqMan MicroRNA Reverse Transcription kit (Applied Biosystems, Waltham, MA, USA). TaqMan primers were used for *Gapdh*, *Bdnf*, and *Pyroxd2* in TaqMan gene expression assays (Applied Biosystems). qPCR for miR137, miR-132 and snoRNA202 were performed using the TaqMan MicroRNA expression assays (Applied Biosystems). For relative quantification of mRNA expression (*Bdnf*, *Pyroxd2*), Arithmetic means were calculated using the comparative 2−ΔΔCt method, with the housekeeping gene *Gapdh* used as the endogenous reference. snoRNA202 was used as the endogenous reference genes for miR-132 and miR-137. Each sample was assayed in triplicate. **All primers are listed in the table below.**

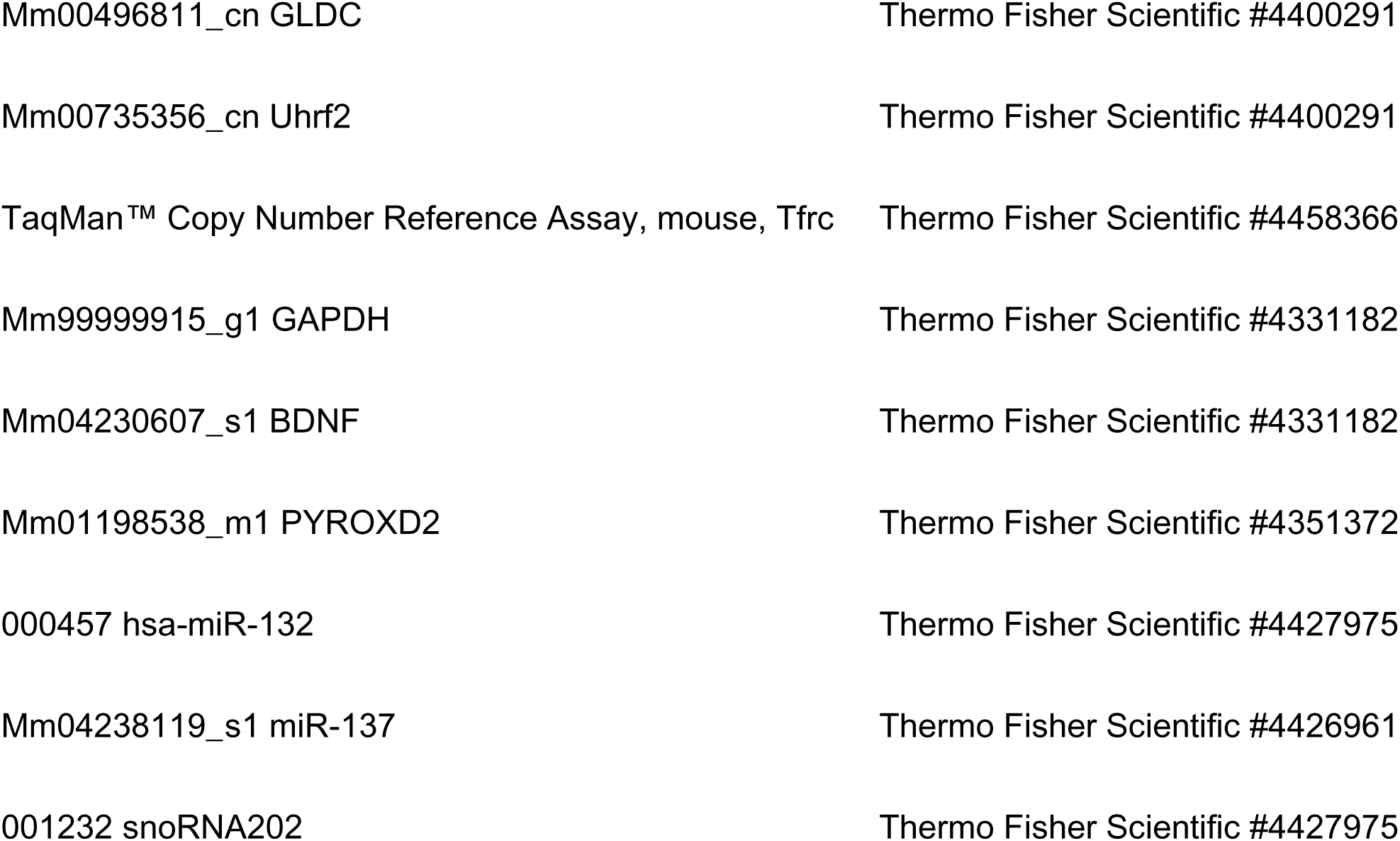

### Immunofluorescence, confocal imaging and image analysis

Mice were deeply anesthetized with Ketamine (Covetrus, OH) / Xylazine (AKORN animal health, IL) (139 mg/kg / 21 mg/kg i.p.) and were perfused transcardially with ice-cold phosphate-buffered saline (PBS), followed by ice-cold 4% paraformaldehyde. The brains were dissected and post-fixed in the same fixation solution for 24 hr and transferred to 30% sucrose solution for cryoprotection for about three days. The brains were sectioned coronally into 40 um-thick sections using cryotome and collected as floating sections in PBS. A few sections were then processed for antigen retrieval.

Sections were taken into the 24-well plate and covered with aluminum foil. This plate was placed in a pre-heated 10mM sodium citrate buffer for 5 min at 90°C in water bath for antigen retrieval. After 5 min of treatment, the plate was kept at room temperature (15 min) to allow the cooling of the buffer. The sections were transferred to the new 24-well plate and three washes with PBS for 6 min each. Then the sections were subjected to the blocking reagent (2% NGS (normal goat serum) + 0.4% Triton X-100 in PBS 1X) for 2 hours at room temperature. With no wash step, the sections were subjected to the primary antibodies in blocking solution, stored at 4°C (cold room) for 2 days. The sections were washed three times with blocking solution for 6 min each and the secondary antibody solution is added in blocking solution for 2 hours at room temperature. Followed by three washes with 1X PBS + 0.05% Tween 20 for 6 min each. DAPI staining was performed by placing the sections in the DAPI (200X) solution for 10 min and sections were taken on to the slides and allowed to dry. Coverslip was mounted with Vectashield antifade Mounting Medium (VectorLabs, Newark, CA, USA).

All brain sections were visualized with (Nikon Eclipse Ti) confocal microscope equipped with Lumencor’s SOLA Light Engines modern solid-state illumination. Images were scanned with ×10, ×20 and ×40 objectives for analysis and qualitative investigation with NIS Elements imaging software. Three to four sections consisting of fully intact and undamaged hippocampal sub-regions were used for image acquisition. Using different channels to visualize different target proteins, images were acquired and saved in TIFF format. Finally, the ImageJ software was used to merge all the different channels for an integrated view of all the proteins in each target region of hippocampus. **All antibodies are listed below.**

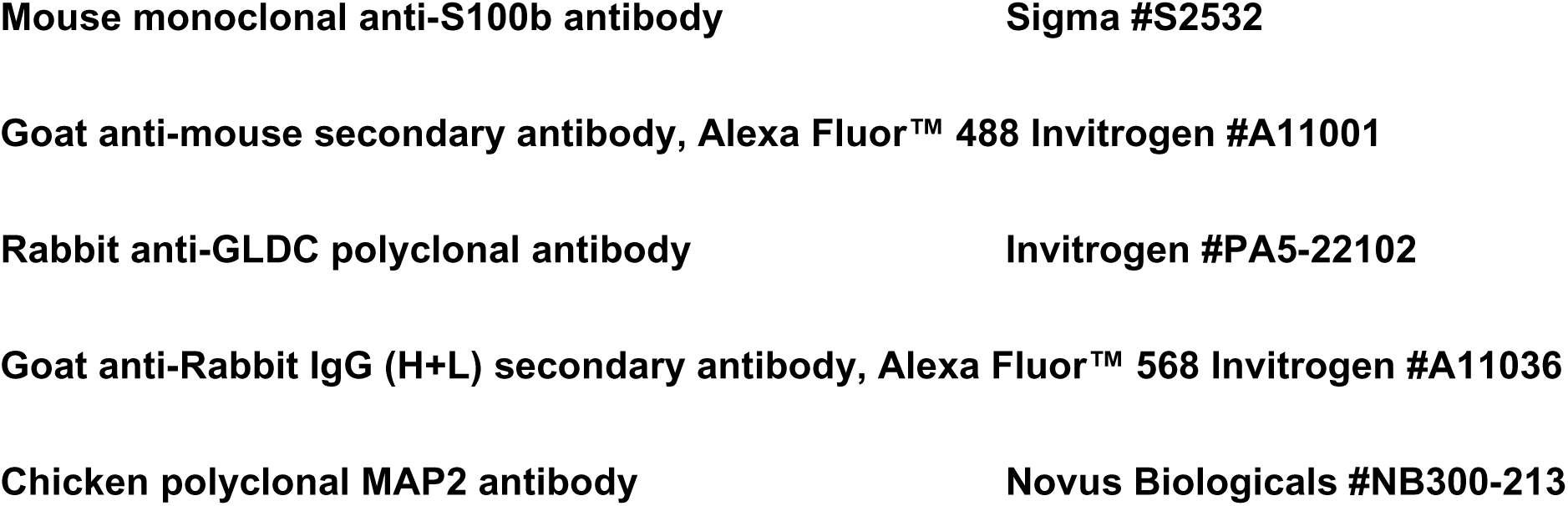
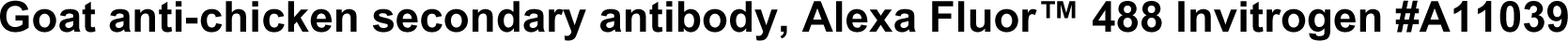

### Data Analysis

Sample sizes were based on established practice in respective assays, and to some extent limited by availability of mice. For behavioral tests, Western blot, qPCR and qRT-PCR, statistical analysis was performed using SigmaPlot 11.0 (Systat Software, USA) or Prism 5 or Prism 7 (GraphPad Software, USA) unless otherwise specified. Results were analyzed using two-sided Student t-test, or one-way ANOVA (when comparing >2 groups) with post-hoc test Bonferroni’s multiple comparisons method or two-way repeated-measures ANOVA, followed by multiple comparisons with Bonferroni’s correction where appropriate (i.e., only when a significant [p < 0.05] main effect was detected). Two-tailed levels of significance were used and for all tests alpha value set at < 0.05.

### RNASequencing and Analysis

#### RNAseq library preparation

##### Strand-specific dUTP RNAseq library preparation

RNA was extracted from the mouse tissues (HPC and mPFC) of deletion, wild type, duplication and triplication mice (n=8 per genotype; 4M, 4F) using TRIzol Reagent. RNA concentration and quality was assessed by TapeStation with RIN ≥8 required for downstream use for RNAseq or qPCR. RNASeq libraries (n=64) were prepared using TruSeq® Stranded mRNA Library Kit (Illumina) and prepared per manufacturer’s instructions. In brief, RNA sample quality (based on RNA Integrity Number, RIN) and quantity were determined on an Agilent 2200 TapeStation and between 500-100 ng of total RNA was used to prepare libraries. 1 uL of diluted (1:100) External RNA Controls Consortium (ERCC) RNA Spike-In Mix (Thermo Fisher) was added to each sample alternating between mix 1 and mix 2 for each well in batch. PolyA bead capture was used to enrich for mRNA, followed by stranded reverse transcription and chemical shearing to make appropriate stranded cDNA inserts for the library. Libraries were finished by adding both sample-specific barcodes and adapters for Illumina sequencing followed by between 10-15 rounds of PCR amplification. Final concentration and size distribution of libraries were evaluated by 2200 TapeStation and/or qPCR, using Library Quantification Kit (KK4854, Kapa Biosystems), and multiplexed by pooling equimolar amounts of each library prior to sequencing. RNASeq libraries were sequenced to approximately 40 million reads per library. These libraries were sequenced on Illumina HiSeq machines.

##### RNASeq preprocessing

FATSQC was used to determine the data quality of the sequencing data. The read pairs were then aligned to either mouse genome (mm10, GRCm38 Ensembl release 83, n=144) using STAR 2.5.3 ^65^ with following parameter–“--outFilterMultimapNmax– --outFilterMismatchNoverLmax 0– --alignEndsType local”. STAR was also used to quantify read counts mapping per gene based on GTF files. Picard and RNAseqQC were used for quality control of the data. The samples with less than 2M estimated library size (https://broadinstitute.github.io/picard/) were removed. Counts per million (CPM) were calculated based on the number of uniquely mapped reads. 0.1 CPM cut-off in 50% samples of one type of sample in the comparison were used to filter the genes with lower expression. These filtered genes were used for differential and correlation analysis.

##### Differential Gene Expression analysis

Differential expression analysis was performed using R package DESeq2 version 1.16.1^65^. The CPM filtered genes counts were used as input for DESeq analysis. DEG was performed with tissue (HPC, mPFC) and edited samples (dels, dups and trips) were compared to corresponding wild type samples. PCA was used to determine the clustering of samples in all the comparisons.

Tissue was the most significant distinguishing factor in all samples but did not explain all the variability. To estimate the hidden variables, Surrogate variable analysis (SVAseq) was used to only preserve the genotype. The hidden variables estimated using SVA were regressed out of the counts and the sva corrected counts were used for DEG and WGCNA analysis. DESeq2 was used to the estimation of size factors (controlling for differences in the sequencing depth of the samples), the estimation of dispersion values for each gene, and fitting a generalized linear model. This resulted in estimated log2 fold changes and p values, which were corrected for multiple testing using Benjamini-Hochberg adjusted p-values (adj. p-value). The significant DEGs were selected if adj. p-value < 0.1. We tested enrichments of DEGs using one-tailed Fisher’s exact test against neurological diseases associated genes from rare variants association analysis, namely ASD ^20^, NDD ^20^ and SCZ ^66^.

## Data availability

All data are included in the manuscript and/or Supplementary Material. RNA sequencing data are deposited in the NCBI GEO database with the following accession number: GSE230871.

## Code availability

No custom code was used for any part of the data processing or analysis.

## Acknowledgements

UR thanks Dr. Bruce M. Cohen, McLean Hospital and Harvard Medical School, and Dr. Edward M. Scolnick, Broad Institute of Harvard and MIT, for their support of seed funding from the Shervert Frazier Research Institute at McLean Hospital and the Stanley Center for Psychiatric Research at the Broad Institute, respectively, Dr. Herman Wolosker (Technion-Israel Institute of Technology) for helpful discussions, and Kelly Brown (McLean Hospital) for performing water T maze experiments. We thank Drs. Xinzhu Yu, Wenyan Mei and Makoto Inoue (UIUC) for providing antibodies for immunofluorescence.

Research in this paper was further supported by a Harvard Brain Science Initiative Bipolar Disorder Seed Grant, supported by Kent and Liz Dauten, to UR and VYB, and by the National Institute of Mental Health of the National Institutes of Health under award numbers R21MH104505 and R56MH112642 to UR, MH115957, HD096326, MH123155, NS093200 to MET and RY, and P50MH115874; R01MH123993; R01MH108665 to VYB and MH51290 to JTC. The content is solely the responsibility of the authors and does not necessarily represent the official views of the National Institutes of Health.

This paper is dedicated to the memory of Dr. Deborah L. Levy (1950-2020), psychologist and researcher at McLean Hospital and Harvard Medical School, who conducted family studies that were essential for the identification and characterization of the marker chromosome in the two patients and who had a crucial role in initiating the studies reported here.

## Author contributions

This study was conceived, designed and supervised by UR. Generation of genetically modified mouse models by J. Liu, JGC, EE, and GEH. PM performed and analyzed the molecular and behavioral assessment of the full CNVs. MK performed and analyzed experiments for partial complementary CNVs. MK performed and analyzed molecular experiments, immunofluorescence, GLDC enzyme activity assay. MK and MW performed and analyzed mitochondrial bioenergetics assay, neuronal morphological and spine density analysis. MK performed various behavioral experiments and analysis. YL and VYB performed and analyzed electrophysiological experiments, PU, JFP and CH performed and analyzed extracellular glycine measurements, RY, JS and MET performed and analyzed RNA sequencing experiments, VV and CJJ provided GlyFS. MW, RN, J. Lyu analyzed experiments. MK prepared graphs for molecular and behavioral analysis of the full CNVs and graphs for RNAseq results. MK prepared figures, MK, VYB, CH and UR wrote the manuscript. All authors revised and approved the final version of the manuscript.

## Additional information

**Correspondence and requests for materials** should be addressed to Uwe Rudolph.

---

**Extended Data Fig 1.**
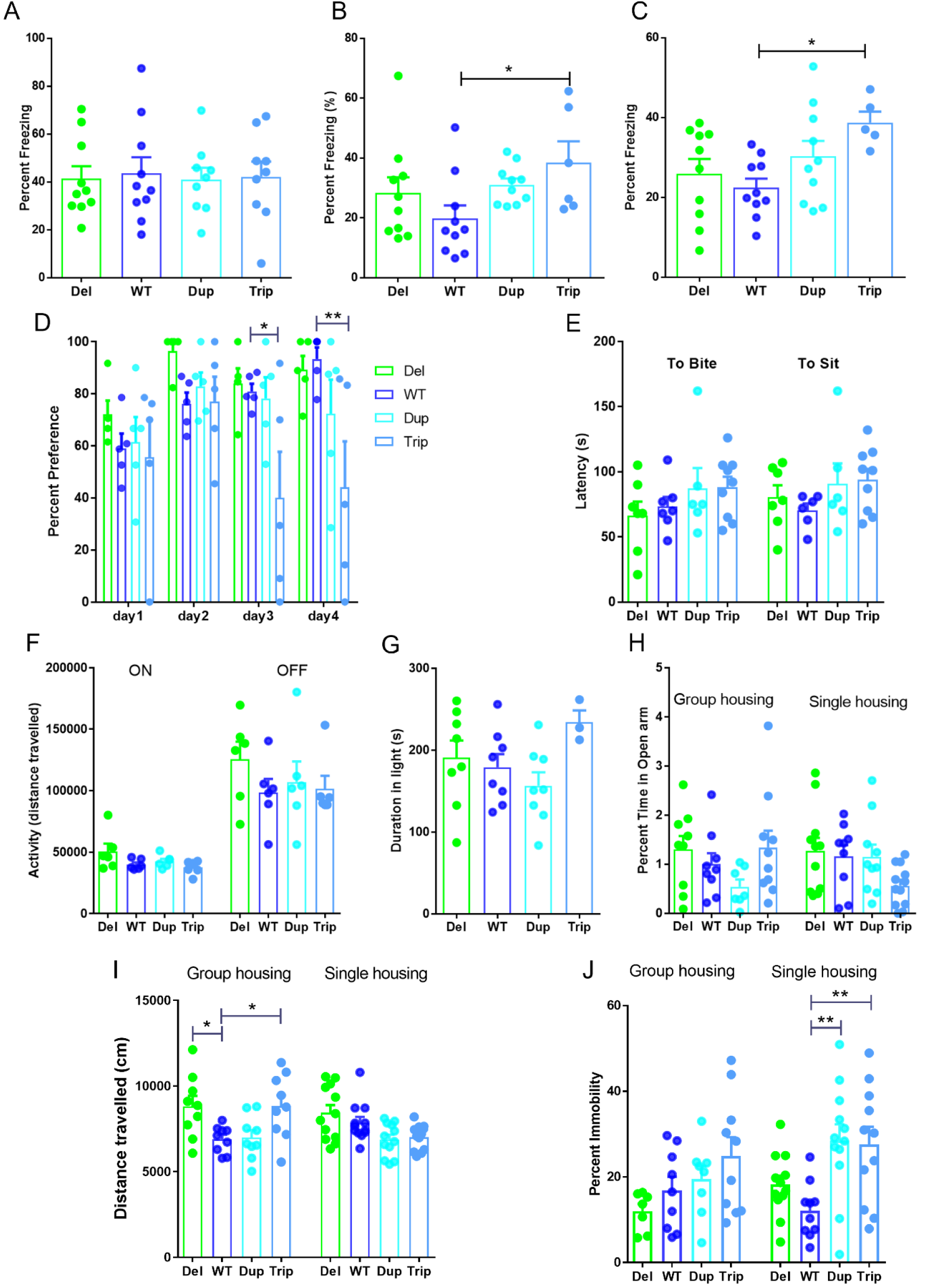
Effect of 9p24.1 CNVs on various cognitive and affective behaviors. A-C, Fear conditioning. In fear conditioning experiments with a shock intensity of 0.5 mA, mice with altered copy numbers of the 9LR segment showed no difference in hippocampal-independent delay conditioning (A), but mice with 4 copies of the 9LR genes (4c9LR mice) displayed increased freezing in hippocampal-dependent trace fear conditioning (B) (One-way ANOVA followed by Bonferroni’s test, F(3,32)=2.493; p=0.0778; 4c9LR vs wildtype, t=2.626, *p<0.05, n=6-10/group) and contextual fear conditioning (C) (One way ANOVA followed by Bonferroni’s test, F(7,63)= 2.343; *p=0.0343; 4c9LR vs wildtype, t=2.468, *p<0.05). D. In the sucrose preference test all genotypes showed a similar preference to sucrose (0.5% w/v vs. water) on days 1 and 2, however, on day 3 and 4 the 9p24.1 triplication mice (4c9R) displayed a reduced preference for sucrose, indicating anhedonia (Two way ANOVA showed significant main effect of days of treatment; F(3,16)=3.374; *p=0.0445 and main effect of genotype; F(3,16)=7.681; ***p=0.0003) followed by Bonferroni’s test, 4c9LR vs wildtype, day3 t=3.020, *p<0.05; day4 t=3.631, **p<0.01, n=5). E. The novelty-suppressed feeding test showed no difference in the latency to bite the pellet and the latency to sit and eat in the chamber compared to WT in any 9p24.1 CNV mice (n=6-10). F. The locomotor activity in an open field chamber was measured separately for the dark (OFF) and light phases (ON) in a 48-hour period, 9p24.1 CNV mice showed similar activity as WT mice in both dark and light phases (n=6). G. In the light/dark choice test, the time spent in the lit chamber was not statistically different from WT in any of the 9p24.1 CNV mice (n=3-8). The following experiments were performed under the two conditions group housing and six weeks of single housing, the latter allowing to assess the effect of social isolation stress. H. In the elevated plus maze test, the percent time in the open arms was not significantly altered in any of the 9p24.1 CNV mice as compared to WT in both conditions (n=7-12). I. In a familiar open field test, group-housed but not single-housed mice with 1 and 4 copies of the 9LR segment (1c9LR=Del, 4c9LR=Trip) displayed an increased distance travelled compared to mice with 2 copies of the 9LR segment (2c9LR=WT) (One way ANOVA, F(7,83)=4.422; ***p=0.0004 followed by Bonferroni’s test, 1c9LR vs wildtype; t=3.085, *p<0.05, 4c9LR vs wildtype, t=3.126, *p<0.05, n=8-12). J. In the forced swim test (6 min), in group-housed mice 9LR copy number had no effect on the time immobile (“1.0” equivalent to 100%), whereas in single-housed mice, mice with 3 or 4 copies of the 9LR segment (3c9LR=Dup and 4c9LR=Trip) displayed an increase in the time spent immobile and thus behavioral despair (One way ANOVA, F(3,44)=5.774; **p=0.0022 followed by Bonferroni’s test, 3c9LR vs wildtype; t=3.558, **p<0.01, 4c9LR vs wildtype, t=3.294, **p<0.01, n=7-12) (J). All data presented as mean ± SEM.

**Extended Data Fig 2.**
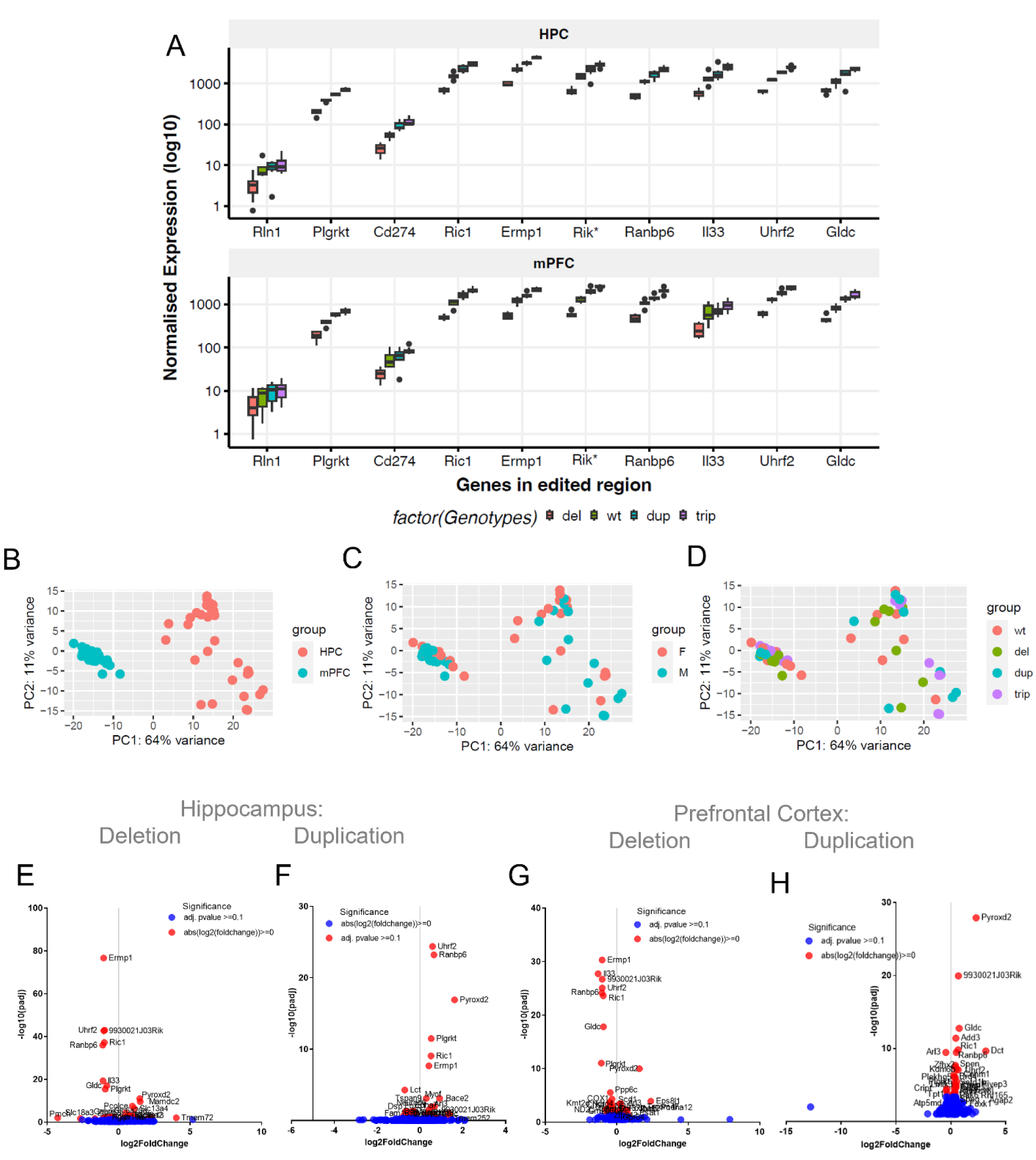
Expression profiling of genes in 9p24.1 cytoband in hippocampal and prefrontal cortex, and differentially expressed genes in duplication and deletion models. A. The box plots show normalized expression of genes in the region of 9p24.1 cytoband with deletion, duplication, and triplication modifications in both hippocampus (top panel, HPC) and prefrontal cortex (bottom panel, mPFC). The x-axis as the order of genes in 9p24.1 cytoband and the y-axis the normalized expression in Log_10_ scale. The gene Rik* = 9930021J03Rik. B-D. The PCA plots of the samples based on top 500 genes with highest variations, tissue level-based variations (B), Gender level-based variations (C) and genotype-based variations (D). This reveals the point that tissue-level variation is the strongest variation among the known batch variables. E-H. The volcano plot shows the differentially expressed genes in hippocampus for (E) deletion and (F) duplication mice, and in prefrontal cortex of (G) deletion and (H) duplication mice. The volcano Plot with the x-axis shows the log2 fold-change and the y-axis as the negative log_10_ of the adjusted p-values. Downregulated genes are shown in the left quadrant and upregulated genes are shown in the right quadrant with top 30 differentially expressed genes (DEGs) in red and others in blue.

**Extended Data Fig 3.**
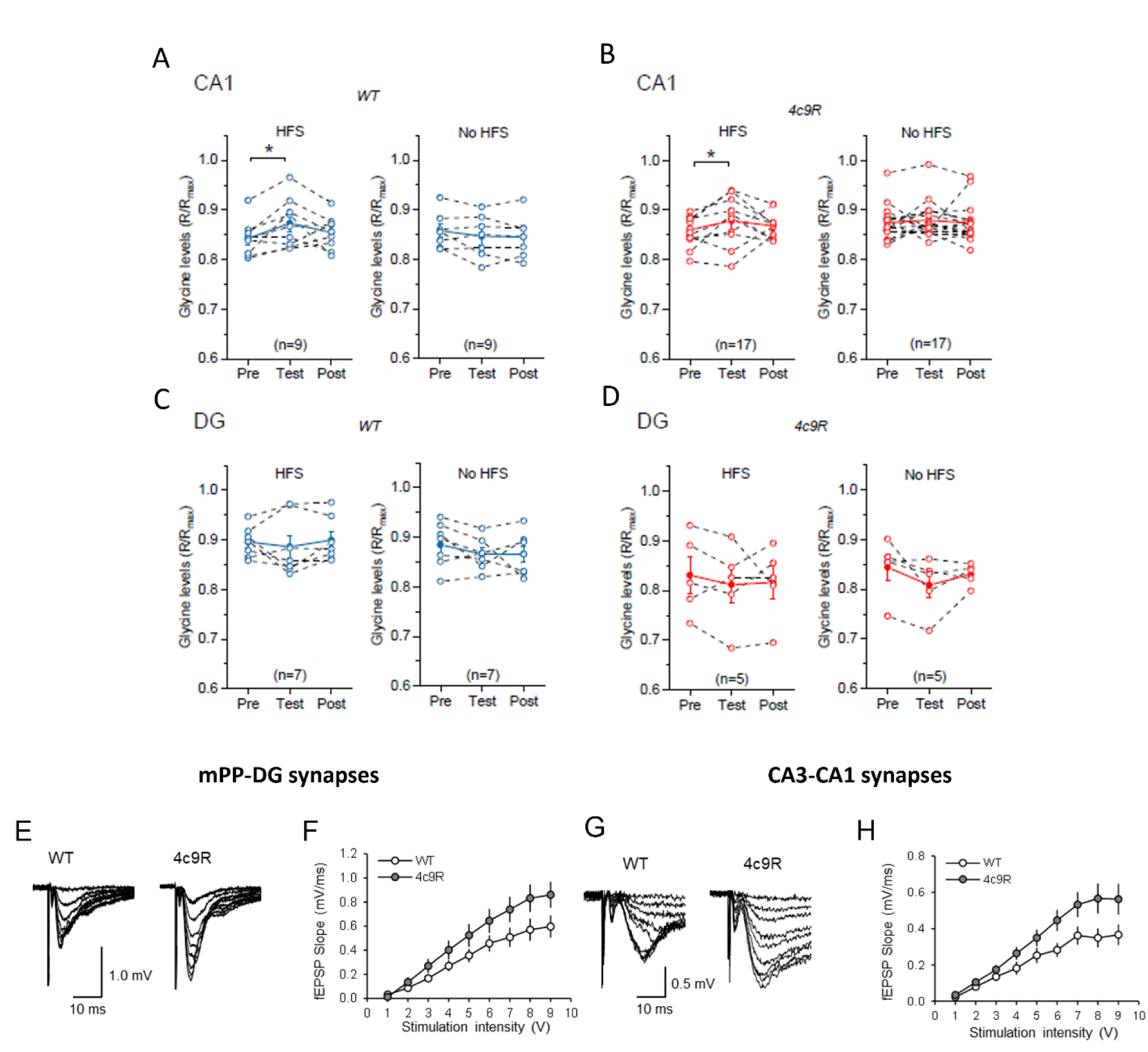
Extracellular glycine concentrations after high frequency stimulation (HFS) in CA1 and DG of 4c9R mice. A-D. High-frequency stimulation (HFS) of Schaffer collateral CA3-CA1 synapses (A, B) and of perforant pathway-DG synapses (C, D) while monitoring GlyFS fluorescence. The pre-test baseline, test response and a post-test baseline were determined. A significant increase of extracellular glycine levels was detected in CA1 from WT mice (paired two-sided Student t-test, t(8) =-3.13, *p=0.014; n= 9 independent experiments) (A, left panel) and 4c9R mice (paired two-sided Student’s t-test, t(16) =-2.66, *p=0.017; n= 17 independent experiments) (B, left panel) and no change was seen when HFS was omitted in CA1 of both WT (paired two-sided Student’s t-test, t(8) = -0.35, p=0.74; n= 9 independent experiments) (A, right panel) and in 4c9R (paired samples Wilcoxon signed ranks test, Z =-0.66, p=0.51; n= 17 independent experiments) (B, right panel). In the dentate gyrus, no increase of GlyFS-reported glycine levels was observed with HFS in WT mice (paired samples Wilcoxon signed ranks test, Z=0.42, p=0.67; n= 7 independent experiments) (C, left panel) and 4c9R mice (paired two-sided Student’s t-test, t(4) =1.16, p=0.31; n= 5 independent experiments) (D, left panel). Without HFS there was no change in WT mice (paired two-sided Student’s t-test, t(6) =1.81, p=0.12; n= 7 independent experiments) (C, right panel) and 4c9R (paired two-sided Student’s t-test, t(4) =1.97, p=0.12; n= 5 independent experiments) (D, right panel). Individual experiments are shown as dots; data are presented as mean ± SEM in blank bars where applicable. All data presented as mean ± SEM, 4 male+ 4 female WT mice and 5 male + 3 female 4c9R mice. We found that there was a significant increase in magnitude of response with increasing input stimuli during stimulus response test at both mPP-DG and CA3-CA1 synapses, but the averaged paired-pulse ratio was unchanged at both synapses. *Averaged fEPSPs (5 traces) evoked at E. mPP-DG synapses and G. CA3-CA1 synapses by presynaptic stimuli of increasing intensity in slices from WT (Left) and 4c9R mice (Right). Synaptic input-output curves of fEPSPs recorded in slices from WT mice (n = 7 slices) and in slices from 4c9R mice (n = 7 slices). Two-way ANOVA for F. mPP-DG synapses: F(1,108) = 18.659, P < 0.001; and H. for CA3-CA1 synapses: F*_(1,135)_ = 20.841, *P* < 0.001.

**Extended Data Fig 4.**
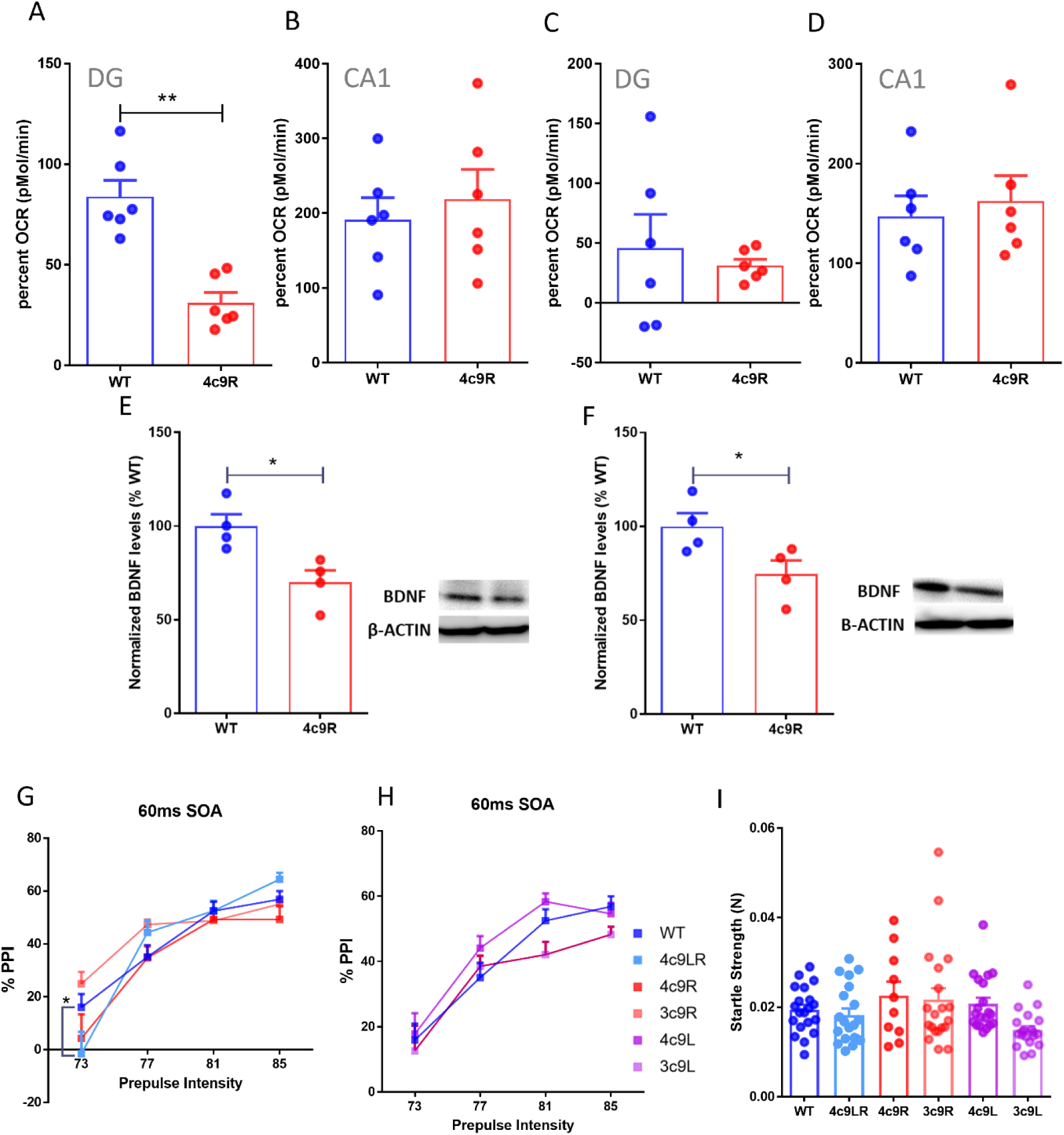
Mitochondrial bioenergetics and BDNF protein levels in hippocampal subregions, and functional genomic fine mapping of prepulse inhibition. A-D. Seahorse mitochondrial stress test (different presentation of the data presented in Fig. 6F and 6H). Maximal respiration was significantly reduced in 4c9R mice compared to wild type in DG (A) (unpaired two-sided student t-test; t(4)=5.370, **p<0.01) but not altered in CA1 (B). Spare/reserve capacity was not altered in DG (C) or CA1 (D). All data presented as mean ± SEM from six mice (4 males and 2 females) per group. E, F. Hippocampal subregional expression of neurotrophic factor BDNF in CA1 and dentate gyrus of 9c4R mice. BDNF protein expression was decreased in 4c9R mice in both CA1 (E) (unpaired two-sided student t-test; 4c9R vs WT (t(6)=3.325; *p<0.05) and DG (F) (unpaired two-sided student t-test; WT vs 4c9R (t(6)=2.502; *p<0.05) compared to WT (n=4, 3 male and 1 female mice per group). G, H. A prepulse inhibition test using different prepulse intensities with 60ms stimulus onset asynchrony (SOA) using WT, 4c9LR, 4c9R, 3c9R mice (G), two-way repeated measures ANOVA showed significant main effect of prepulse intensity (F(3,85) =110.3, ***p<0.001) on PPI and a significant interaction between prepulse intensity and genotype (F(15,85) =1.954, *p<0.05). Further, the Bonferroni test showed a significant difference between 4c9LR vs WT at 73dB prepulse intensity at 60ms SOA (t= 2.599, *p<0.05) (G) but not in 4c9R, 3c9R, 4c9L, 3c9L mice (H). I. Responses to no stimulus trials during the prepulse inhibition test were not altered in any genotypes compared to wild type. n= 10 with 5 male and 5 female mice. All data presented as mean ± SEM.

**Extended Data Fig 5.**
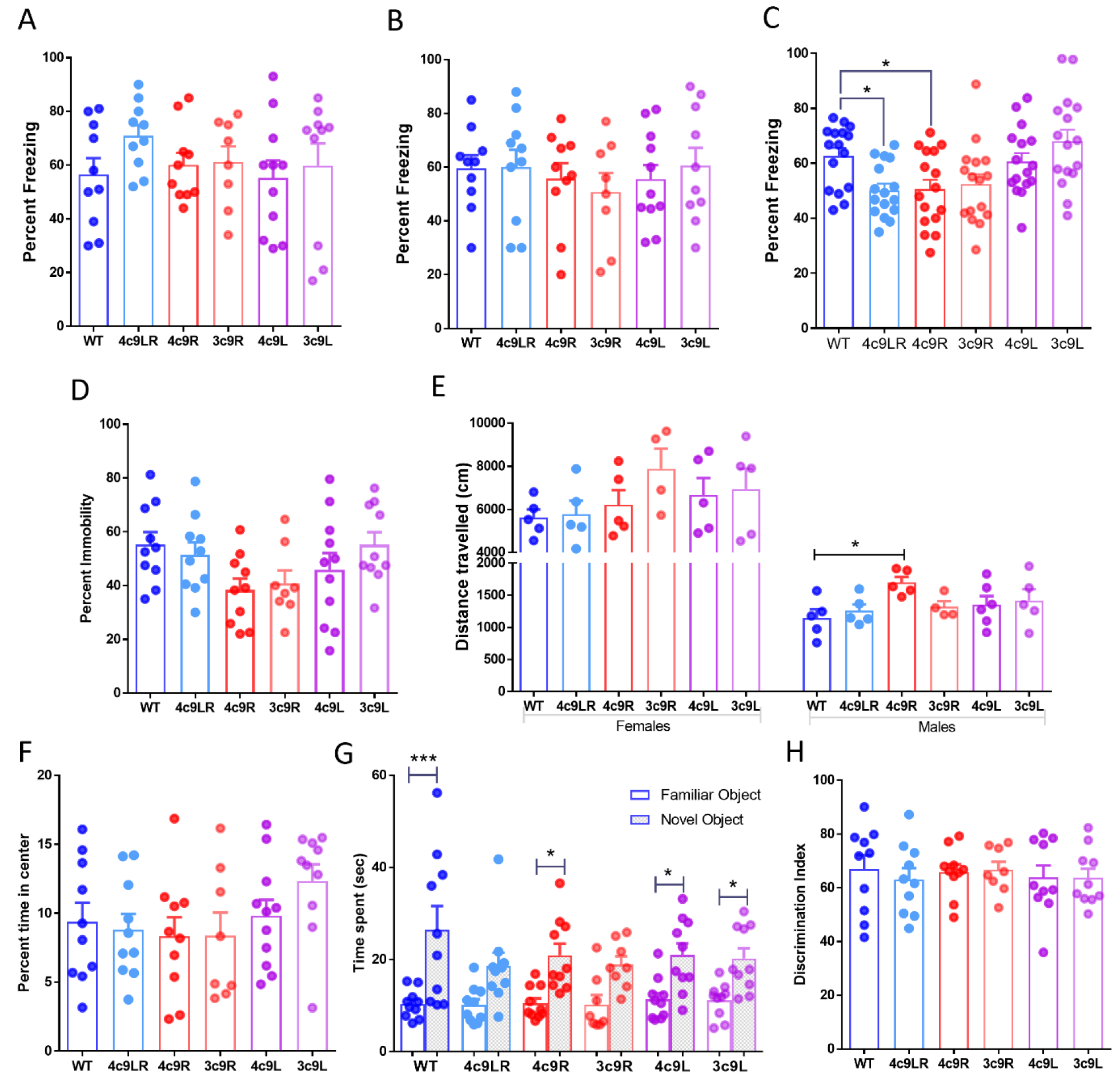
Functional genomic fine mapping of various cognitive and affective behaviors. A-C. Fear conditioning. Hippocampal-dependent trace fear conditioning with 0.7 mA shock intensity (A) and contextual fear conditioning (B) did not alter the freezing response in any of the genotypes 4c9RL, 4c9R, 3c9R, 4c9L and 3c9L compared to WT (n=8-10). C. Delay fear conditioning with 0.7 mA shock intensity, representing the non-preexposed data from the latent inhibition experiment. The mice that contain 4 copies of GLDC (4c9LR and 4c9R) show reduced freezing response (One way ANOVA, F(5,90)=4.957; ***p=0.0005 followed by Dunnett test, 4c9LR vs wildtype; q=2.667, *p<0.05, 4c9R vs wildtype; q=2.585, *p<0.05, n=16, 8 males and 8 females). D. In the forced swim test (10 min), the percentage of time spent immobile was not altered for any of the genotypes 4c9RL, 4c9R, 3c9R, 4c9L and 3c9L compared to WT (n=8-10). E. In the open field test, total distance travelled was increased in male 4c9R mice (One way ANOVA, F(5,24)=2.120; p=0.0977 followed by Bonferroni’s test, 4c9R vs wildtype; t=3.039, *p<0.05, n=8-10), but not in any other genotype (E, right panel), and not in females (E, left panel). F. The percent time spent in the center was not altered in any of the genotypes 4c9RL, 4c9R, 3c9R, 4c9L and 3c9L compared to WT (n=8-10). G, H. In the novel object recognition test, WT, 4c9R, 4c9L and 3c9L mice which showed a preference exploring the novel object compared to familiar object, while 4c9LR and 3c9R mice did not exhibit such a difference (One way ANOVA, F(11,99)=5.694; ***p<0.0001 followed by Bonferroni’s test, WT; t=4.747, ***p<0.001; 4c9R; t=3.077, *p<0.05, 4c9L; t=2.846, *p<0.05, 3c9L; t=2.718, *p<0.05, n=8-10) (G). However, the discrimination index showed no genotypic differences (H). All data presented as mean ± SEM from 4-5 males+ 4-5 female mice each group.

